# Recombination-independent recognition of DNA homology for meiotic silencing in *Neurospora crassa*

**DOI:** 10.1101/573907

**Authors:** Nicholas Rhoades, Tinh-Suong Nguyen, Guillaume Witz, Germano Cecere, Thomas Hammond, Alexey K. Mazur, Eugene Gladyshev

**Author notes:** Corresponding author: E.G.

## Abstract

Pairing of homologous chromosomes represents a critical step of meiosis in nearly all sexually reproducing species. While in some organisms meiotic pairing requires programmed DNA breakage and recombination, in many others it engages homologous chromosomes that remain apparently intact. The mechanistic nature of such recombination-independent pairing represents a fundamental question in molecular genetics. Using ‘meiotic silencing by unpaired DNA’ (MSUD) in *Neurospora crassa* as a model process, we demonstrate the existence of a cardinally different approach to DNA homology recognition in meiosis. The main advantage of MSUD over other experimental systems lies in its ability to identify any relatively short DNA fragment lacking a homologous allelic partner. Here we show that MSUD does not rely on the canonical mechanism of meiotic recombination, yet it is promoted by REC8, a conserved component of the meiotic cohesin complex. We also show that certain patterns of interspersed homology are recognized as pairable during MSUD. Such patterns need to be co-linear and must contain short tracts of sequence identity spaced apart with a periodicity of 21 or 22 base-pairs. By using these values as a guiding parameter in all-atom molecular modeling, we discover that homologous double-stranded DNA molecules can associate by forming quadruplex-based contacts with an interval of 2.5 helical turns, which requires right-handed plectonemic coiling and additional conformational changes in the intervening double-helical segments. These results (i) reconcile genetic and biophysical lines of evidence for the existence of direct homologous dsDNA-dsDNA pairing, (ii) identify a role for this process in initiating post-transcriptional silencing, and (iii) suggest that chromosomes are cross-matched in meiosis by a precise mechanism that operates on intact double-stranded DNA molecules.

## INTRODUCTION

Correct pairing of homologous chromosomes in meiosis is essential for ensuring fertility and preventing birth defects. While in some organisms this process depends on programmed DNA breakage and recombination, in many others it engages apparently intact chromosomes (1). Such recombination-independent pairing was proposed to rely on indirect sequence readouts by locus-specific proteins or RNAs, while also being guided by landmark chromosomal features, such transcriptional hubs, heterochromatin domains, centromeres, and telomeres (2–6). Alternatively, a possibility exists that recombination-independent pairing can be established at the DNA level (7, 8), but a mechanistic model, consistent with *in vivo* observations, has been lacking (9).

A process known as ‘meiotic silencing by unpaired DNA’ (MSUD) induces transient RNA interference against heterologous DNA sequences found at the allelic chromosomal positions during meiosis (10). In the fungus *Neurospora crassa*, the ability of MSUD to detect unpairable DNA segments as short as 1 kbp suggests that an efficient homology recognition mechanism is involved (11). This mechanism cannot rely on any premeiotic pairing, since meiosis occurs immediately after the fusion of haploid parental nuclei in *N. crassa* and related fungi. More than two dozen MSUD factors have been identified with roles in transcription, mRNA processing, RNA interference, chromatin remodeling and nuclear dynamics, but its mechanism of homology recognition remained unknown (11, 12).

In addition to MSUD, *N. crassa* has another homology-dependent process known as ‘repeat-induced point mutation’ (RIP), which takes place exclusively in premeiotic haploid nuclei (13, 14). During RIP, gene-sized duplications of genomic DNA become mutated by numerous cytosine-to-thymine transitions along their entire lengths (13, 14). The capacity of RIP to detect DNA repeats irrespective of their particular sequence, coding potential and genomic positions suggests that it involves an exhaustive genome-wide homology search. Yet RIP still functions in the absence of MEI-3 (the only RecA-like recombinase in Neurospora) and SPO11 (a topoisomerase-like protein that makes DNA breaks to initiate meiotic recombination) (15). By analyzing the ability of synthetic repeats with different patterns of homology to induce mutations, it was discovered that RIP could detect the presence of matching trinucleotides (triplets), but only when they were interspersed with a periodicity of 11 or 12 bps between the two participating DNA segments (15). Taken together, these results uncovered the existence of a cardinally different mechanism of DNA homology search and recognition that likely cross-matches intact double-stranded DNA molecules directly, without involving single-stranded DNA (or RNA) intermediates (15).

Recognition of interspersed homology for RIP was accounted for by a recently proposed all-atom model of direct dsDNA-dsDNA pairing (16). This model is based on the physical property of standard Watson-Crick (WC) base-pairs to present distinct yet self-complementary electrostatic patterns along their major-groove edges. This property allows homologous double-stranded stacks to pair without breaking WC bonds (16). According to the model, a sequence-specific contact between two DNA double helices represents a short stack of 3-4 planar quartets formed by identical (homologous) WC base-pairs. The favorable energy of this reaction includes a large non-specific contribution of ionic interactions and a sequence-specific hydrogen bonding term. While the model has defined the biophysical properties of the quadruplex-based contacts, it has not explored the role of intervening double-helical segments in enabling the association of long dsDNA molecules (16).

We now report that MSUD does not depend on the canonical mechanism of meiotic recombination. MSUD also proceeds normally in the absence of heterochromatin formation involving H3K9me3 and H3K27me3. At the same time, MSUD is promoted by REC8, thus revealing a recombination-independent role for this central meiotic factor. Further, we show that MSUD can discriminate between the physical absence of DNA and the absence of DNA homology at the allelic position. Strikingly, we find that interspersed homologies containing short tracts of sequence identity arrayed with a periodicity of 21 or 22 base-pairs are readily recognized as pairable during MSUD. Using this parameter in all-atom molecular modeling, we discover that homologous dsDNAs can associate by establishing quadruplex-based contacts with an interval of 2.5 helical turns. This process requires right-handed plectonemic coiling and additional conformational changes in the intervening double-helical segments. Taken together, our results (i) reconcile genetic and biophysical evidence for the existence of the direct homologous dsDNA-dsDNA pairing, (ii) identify a role for this process in initiating post-transcriptional silencing, and (iii) suggest that meiotic chromosomes can be cross-matched by a mechanism that operates on intact double-stranded DNA molecules.

## RESULTS

In *N. crassa* and related fungi, the meiocyte (ascus) starts to develop shortly after karyogamy (Fig. 1A). The ascus enlarges dramatically during prophase I, which is also when the pairing of homologous chromosomes takes place (Fig. 1A). Each fruiting body (perithecium) contains 100─200 asci at different stages of meiosis. When dissected from the fruiting body, the asci form a rosette surrounded by a mass of auxiliary perithecial cells (Fig. 1B).

**Figure 1.**
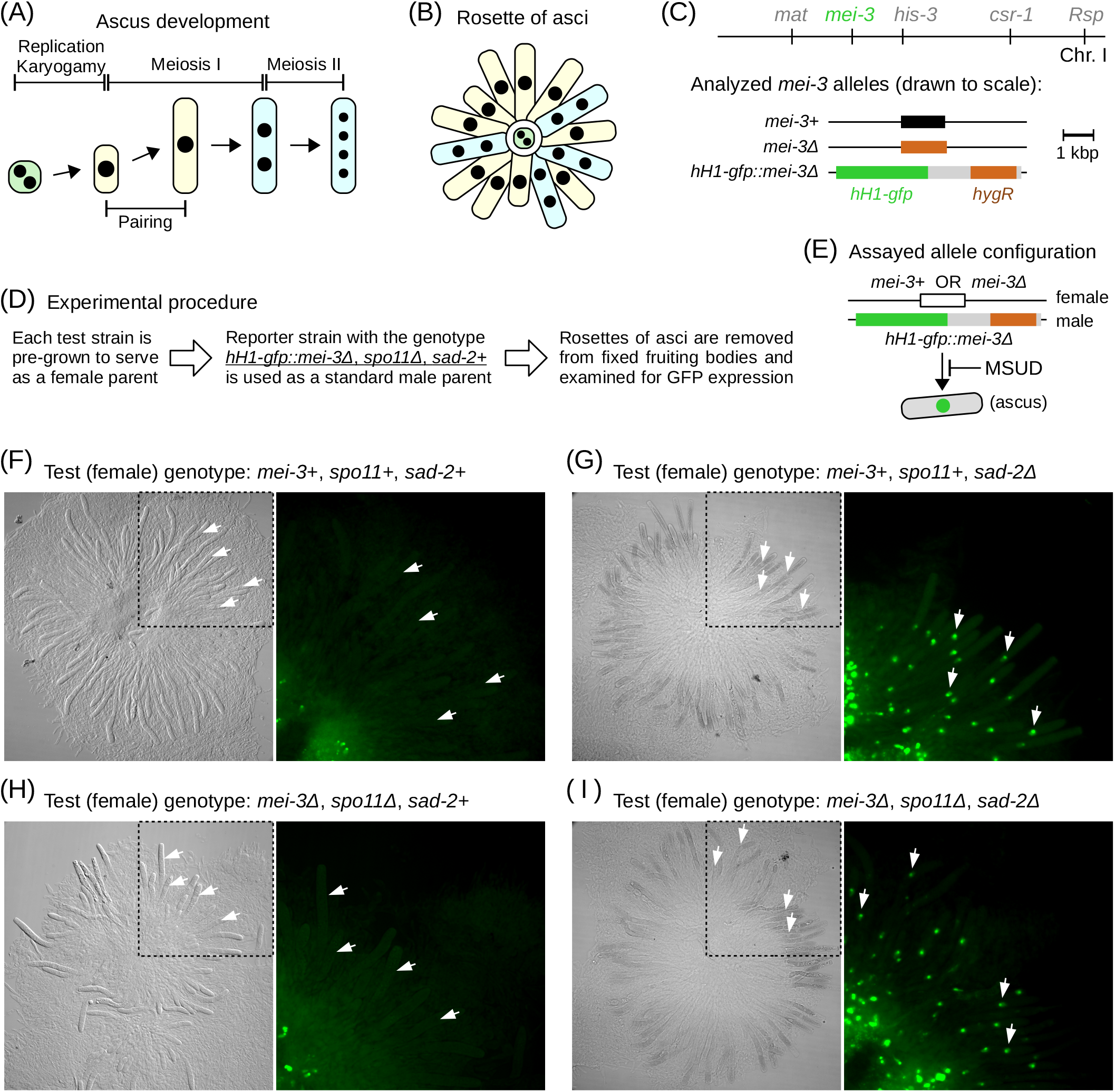
MSUD does not depend on the canonical mechanism of meiotic recombination. (A) In *N. crassa*, meiosis starts immediately after karyogamy. During prophase I, the meiocyte (called ‘ascus’) elongates dramatically and becomes readily recognized. (B) A cartoon representation of a dissected rosette of asci. (C) The reporter construct includes *hH1-gfp* (green) and a hygromycin resistance marker *hygR* (orange). The wildtype *mei-3+* allele was replaced with the construct. The standard deletion allele *mei-3Δ* was produced by the Neurospora Genome Project (70) and obtained from the Fungal Genetics Stock Center (71). All DNA segments are drawn to scale. (D) Overall experimental procedure. GFP_Reporter1 (SI Table 2) is always used as a male parent. (E) The reporter construct remains unpaired because the female test strains provide either *mei-3+* or *mei-3Δ*. (F) Recombination-proficient, MSUD-proficient cross (Cross X1, SI Table 3). (G) Recombination-proficient, MSUD-deficient cross (Cross X2, SI Table 3). (H) Recombination-deficient, MSUD-proficient cross (Cross X3, SI Table 3). (I) Recombination-deficient, MSUD-deficient cross (Cross X4, SI Table 3). Representative meiotic nuclei are indicated with white arrows. All panels showing GFP signal are magnified two-fold, as indicated.

### MSUD does not depend on the canonical mechanism of meiotic recombination

The canonical meiotic pairing program requires SPO11 to produce double-strand DNA breaks (DSBs) and the eukaryotic RecA-like recombinases DMC1 and RAD51 to mediate a subsequent homology search (17). *N. crassa* has only one RecA protein, MEI-3, which is dispensable during vegetative growth but is essential during meiosis (18). Its lethal phenotype, however, can be suppressed by removing SPO11 (15). The ability of *N. crassa* to complete meiosis in the absence of MEI-3 and SPO11 allowed us to test the role of meiotic recombination in MSUD.

A microscopy-based approach was chosen to analyze MSUD by assaying expression of a histone H1-GFP fusion protein (Fig. 1E). Normally expressed hH1-GFP features readily detectable nuclear localization, but it rapidly and completely disappears from meiotic nuclei in heterozygous *hH1-gfp* crosses due to MSUD (19). Furthermore, when the *hH1-gfp* gene is provided solely by the male parent, hH1-GFP remains restricted to premeiotic, meiotic, and postmeiotic tissues, permitting its straightforward detection in dissected rosettes.

In the first reporter strain (GFP_Reporter1), *hH1-gfp* was inserted as a replacement of the *mei-3+* allele (Fig. 1C). GFP_Reporter1 had also the *spo11+* allele deleted. By the overall design, this strain was always used as a standard male parent, while meiotic genotypes were manipulated by using different female strains (Fig. 1D). Depending on which female strain was chosen, crosses could differ with respect to both recombination and MSUD proficiency. The cross could be recombination-proficient (*mei-3+/ΔΔ*, *spo11+/ΔΔ*) or recombination-deficient (*mei-3Δ/ΔΔ*, *spo11Δ/ΔΔ*). Additionally, the cross could be either MSUD-proficient (*sad-2+/Δ+*) or MSUD-deficient (*sad-2Δ/Δ+*). The MSUD deficiency of *sad-2Δ/Δ+* crosses resulted from an effect called ‘silencing the silencer’, in which an unpaired MSUD gene (in this case, *sad-2+*) was subjected to MSUD itself (11).

We found that hH1-GFP was completely and specifically silenced during meiosis in MSUD-proficient crosses (Fig. 1F,H). Critically, the levels of silencing appeared indistinguishable between the recombination-proficient and the recombination-deficient conditions (Fig. 1F,H). Under both regimes, the expression of hH1-GFP was restored by suppressing MSUD (Fig. 1G,I; Fig. S1B,C). The meiotic expression of hH1-GFP remained strong in the recombination-deficient cross with active MSUD if two identical *hH1-gfp* alleles were provided, in other words, if *hH1-gfp* could be paired (Fig. S1A). Together, these results suggest that MSUD does not depend on the canonical mechanism of homologous meiotic recombination involving SPO11 and MEI-3.

### MSUD is not coupled to non-canonical genetic recombination

While SPO11 is needed for chromosome synapsis during meiosis in *N. crassa*, residual SPO11-independent meiotic recombination has been described in this organism (20). Those observations raised a possibility that the wildtype efficiency of MSUD in our *spo11Δ/ΔΔ* crosses could be caused by a SPO11-independent process involving DNA breaks and recombination. To investigate this possibility, we measured genetic recombination over a large interval spanning more than half of chromosome I (Fig. S2A). Notably, this interval included the *his-3* locus, which is characterized by high levels of SPO11-independent recombination in crosses between certain Neurospora strains from a different genetic background (20, 21).

In *N. crassa*, SPO11-deficient crosses produce substantial numbers of aneuploid (partially diploid) progeny (20). These aneuploid progeny may undergo mitotic recombination and chromosome loss during vegetative growth, thus obscuring meiotic outcomes. To minimize the chance of encountering such progeny, *csr-1* was used as one of two reference loci. More specifically, this gene was chosen because (i) only loss-of-function *csr-1* alleles confer resistance to cyclosporin A, (ii) *csr-1*-based resistance to cyclosporin A is recessive, and (iii) *csr-1+* itself is stable during the sexual phase (22). Overall, a recombination-deficient *csr-1+/ΔΔ* cross can produce three *csr-1* genotypes: haploid *csr-1+* and *csr-1Δ* as well as partially diploid *csr-1+/ΔΔ*; however, only *csr-1Δ* progeny should be retained on cyclosporin A. These isolates can be genotyped at the *mat* locus using a PCR-based assay designed to detect a potential mixture of *mat A* and *mat a* (Fig. S2B,C). The presence of *mat A* alleles among the *csr-1Δ* progeny would indicate a genetic exchange had occurred between *csr-1* and *mat* loci. Preliminary analysis confirmed that the overall strategy was effective (Fig. S2D).

We analyzed progeny from two of the crosses that were assayed for *hH1-gfp* silencing (Fig. 1F,H). In these crosses, GFP_Reporter1 provided the *csr-1Δ* allele, while the test strains provided the wildtype *csr-1+* allele. The recombination-proficient cross (Fig. 1F) yielded many recombinants: 26% of the *csr-1Δ* progeny carried the *mat A* allele (Fig. S2E, ‘*mei-3+/ΔΔ*, *spo11+/ΔΔ*’). In contrast, all 594 *csr-1Δ* isolates from the recombination-deficient cross (Fig. 1H) carried the *mat a* allele (Fig. S2E, ‘*mei-3Δ/ΔΔ*, *spo11Δ/ΔΔ*’).

We also considered the possibility that SPO11-independent recombinants were excluded from the offspring of the recombination-deficient cross because they required MEI-3 to complete DNA repair. To this end, we analyzed one additional cross between GFP_Reporter1 and another *spo11Δ* strain that carried the wildtype *mei-3+* allele. From this cross, one recombinant isolate was recovered among the 594 *csr-1Δ* progeny (Fig. S2E, ‘*mei-3+/ΔΔ*, *spo11Δ/ΔΔ*’), suggesting that the level of residual SPO11-independent recombination was low. Taken together, these results show that MSUD is not coupled to genetic recombination.

### MSUD is promoted by REC8

MSUD likely relies on favored interactions between homologous chromosomes rather than sister chromatids. This phenomenon (known as ‘homolog bias’) requires the conserved meiotic kleisin REC8 (23). Neurospora encodes a single REC8 ortholog (NCU03190) with a canonical structure (Fig. S3A). Transcription of *rec8+* is strongly upregulated in perithecia together with other meiotic genes (24).

To evaluate the role of REC8 in MSUD, we created a new reporter (GFP_Reporter2) by integrating *hH1-gfp* as the replacement of *csr-1+* in a strain that had both *rec8+* and *spo11+* deleted. The absence of homology between *hH1-gfp* and *csr-1+* (provided by female parents) was expected to silence *hH1-gfp* (Fig. S3B). One REC8-deficient cross (*rec8Δ/ΔΔ*, *spo11Δ/ΔΔ*) and one REC8-proficient cross (*rec8+/ΔΔ*, *spo11+/ΔΔ*) were tested. Both crosses used GFP_Reporter2 as a male parent. SPO11 was eliminated in the REC8-deficient cross to avoid complications associated with potentially aberrant repair of SPO11-induced DNA breaks.

We found that the REC8-proficient cross contained a variable but small number of asci expressing hH1-GFP (Fig. S3D). This result can be explained by either the ‘silencing the silencer’ phenomenon (above) or a less efficient recognition of the *hH1-gfp::csr-1Δ* allele by MSUD. Strikingly, many more asci expressed hH1-GFP in the REC8-deficient condition, indicating a silencing defect (Fig. S3C). This defect was not complete, and hH1-GFP was lost in some neighboring asci (Fig. S3C). Thus, efficient MSUD is promoted by REC8, but it also occurs, albeit less readily, in the absence of this critical meiotic factor.

### MSUD does not depend on the canonical mechanisms of heterochromatin formation

A process known as ‘meiotic silencing of unsynapsed chromatin’ (MSUC) induces accumulation of repressive chromatin marks (such as di- and trimethylated histone H3 lysine 9, H3K9me2/3) on unpaired chromosomes in animals (25). In *C. elegans*, MSUC requires several conserved RNAi factors including the RNA-dependent RNA polymerase EGO-1 (26). Thus, we explored if MSUD involves a heterochromatin-related component.

In *N. crassa*, constitutive heterochromatin requires the lysine methyltransferase DIM-5, whereas facultative heterochromatin requires the lysine methyltransferase SET-7 (Fig. S4A; ref. 27). DIM-5 and SET-7 mediate all H3K9me3 and H3K27me3 in *N. crassa*, respectively (27). Although *dim-5Δ* strains are largely infertile as females, this defect is suppressed by deleting *set-7+* (28). We thus used the ability of *N. crassa* to complete its sexual phase in the absence of DIM-5 and SET-7 to assay the role of heterochromatin in MSUD. For this test, a third reporter strain (GFP_Reporter3) was engineered by replacing the *csr-1+* allele with *hH1-gfp* in a heterochromatin-deficient (*dim-5Δ, set-7Δ*) background.

We analyzed two heterochromatin-deficient crosses, one of which was MSUD-proficient (Fig. S4B), while the other was MSUD-deficient (Fig. S4C). Overall, dissected rosettes were frequently aberrant (data not shown), and very few elongating asci could be found in both conditions, suggesting that the lack of heterochromatin resulted in a strong developmental defect (Fig. S4B,C). Nevertheless, hH1-GFP was still expressed aptly in premeiotic and early meiotic nuclei (Fig. S4B,C). Importantly, hH1-GFP became silenced specifically in the elongating asci of the MSUD-proficient cross (Fig. S4B,C). These results suggest that meiotic silencing in *N. crassa* does not require heterochromatin formation.

### Developing a new quantitative approach to analyzing the homology requirements of MSUD

Thus far, our results show that MSUD parallels RIP in being independent from meiotic recombination. Earlier we suggested that RIP involves a cardinally different homology recognition mechanism that matches double-stranded DNAs directly, without using single-stranded intermediates (15). This idea stemmed from the fact that RIP mutates certain patterns of interspersed homology with an overall identity of only 36%, substantially below the limit of any sequence recognition process that relies on the annealing of complementary strands (15). Thus, it was important to test if MSUD could also recognize such interspersed homologies as pairable.

To understand the homology requirements of MSUD, a new quantitative approach was developed (Fig. 2A-C). This approach relies on a classical MSUD assay, in which meiotic silencing of the *Roundspore+* (*Rsp+*) gene results in the production of oval-shaped (’silenced’) rather than spindle-shaped (’wildtype’) ascospores (10). Two technical advances included: (i) a sensitive reporter system to test engineered homologies (below), and (ii) an image-processing algorithm to automatically segment and classify large numbers of ascospores (also introduced below).

**Figure 2.**
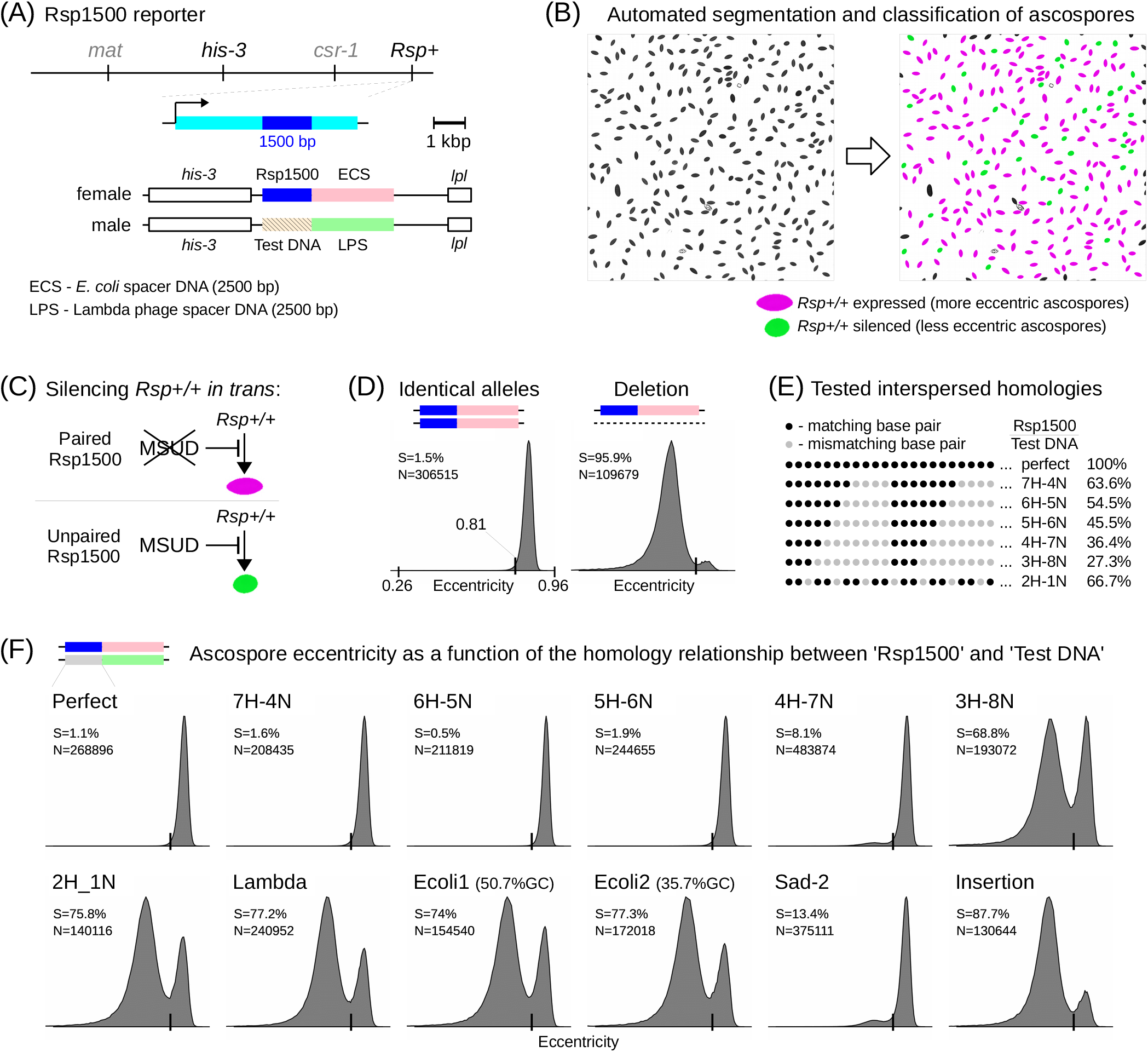
MSUD recognizes weak interspersed DNA homology as pairable. (A) Genetic reporter system to quantify interactions between ‘Rsp1500’ and ‘Test DNA’. (B) Image processing algorithm to classify ascospores. (C) Interactions between Rsp1500 and Test DNA are detected by quantifying the eccentricity of ascospores. (D) The eccentricity threshold of 0.81 is used to classify ascospores as ‘wildtype’ or ‘silenced’. ‘S’ denotes the percentage of ‘silenced’ ascospores (below the threshold). ‘N’ corresponds to the total number of ascospores analyzed for a given condition. Eccentricity distributions are shown as histograms. The following crosses are analyzed: X11 (’Identical alleles’) and X12 (’Deletion’) (SI Table 3). (E) Synthetic Test DNA sequences that form patterns of interspersed homology with Rsp1500 are designed based on the known homology requirements of RIP (15). Overall sequence identities corresponding to each homology pattern are indicated on the right. (F) Eccentricity distributions are displayed as in panel D. Assayed crosses differ only with respect to the Test DNA. The following crosses are analyzed: X13 (’Perfect’), X14 (’7H-4N’), X15 (’6H-5N’), X16 (’5H-6N’), X17 (’4H-7N’), X18 (’3H-8N’), X19 (’2H-1N’), X20 (’Lambda’), X21 (’Ecoli1’), X22 (’Ecoli2’), X23 (’Sad-2’), and X24 (’Insertion’) (SI Table 3).

The reporter system is based on the genetic interaction between ‘Rsp1500’ (an ectopic segment of *Rsp+*) and ‘Test DNA’. Both segments have the same length (1500 bp) and occupy allelic positions near the *his-3* gene (Fig. 2A). While the sequence of Rsp1500 is fixed, the sequence of Test DNA can be varied as desired. Previously, we used a similar approach to dissect the homology requirements of RIP (15).

By overall design, if Rsp1500 and Test DNA are pairable, MSUD will not be induced, the endogenous *Rsp+* alleles will be expressed normally, and spindle-shaped (more eccentric) ascospores will be produced (Fig. 2C). On the other hand, if Rsp1500 and Test DNA are not pairable, MSUD will be activated, the endogenous *Rsp+* alleles will be silenced *in trans*, and oval-shaped (less eccentric) ascospores will be produced (Fig. 2C). To improve the sensitivity of the assay, an adjacent 2500-bp region of heterology was engineered (Fig. 2A). This heterology was formed with two foreign sequences, ‘ECS’ and ‘LPS’ (Fig. 2A). ECS originated in *E. coli*, while LPS was obtained from lambda phage. In this situation, if Rsp1500 and Test DNA are not pairable, the total length of unpaired DNA containing the Rsp1500 segment will be 4 kbp (instead of 1.5 kbp), a condition expected to enhance MSUD (29).

An algorithm was further developed to extract ascospore features from brightfield images (Fig. 2B; Methods). Ascospores are detected using rough localization followed by local segmentation. Segmented objects are filtered based on their convexity and area values, and ascospores that are damaged, aberrant, or clumped together are excluded. The roundness of ascospores is determined using an eccentricity parameter ranging between zero (perfect circle) and one (straight segment) (Fig. 2B,C; Methods). This approach benefited from the fact that large numbers of ascospores could be obtained from crosses in nearly pure form (Fig. 1B).

A reporter strain carrying Rsp1500 adjacent to ECS (RSP_Reporter1) was always used as a female parent (Fig. 2A). Test DNAs were provided by otherwise isogenic strains of the opposite meting type, used as male parents. To calculate the percentage of silenced ascospores, an eccentricity threshold of 0.81 was applied (wildtype ascospores ≥ 0.81). For a given condition, three crosses were analyzed, each represented by 3-15 ×10^4^ ascospores, thus bringing the total number of assayed ascospores to 1-4 ×10^5^.

As a proof of principle, we analyzed two conditions, one featuring two identical reporter alleles and the other featuring a reporter allele and a deletion spanning the entire reporter construct (Fig. 2D). The first condition produced 98.5% wildtype ascospores, while the second condition produced 95.9% silenced ascospores (Fig. 2D). The corresponding distributions of eccentricity values were overlapping but separable. These results show that (i) ascospores can be categorized as wildtype or silenced based on their eccentricities, and (ii) the reporter has a broad dynamic range, enabling a systematic analysis of homology requirements for MSUD.

### MSUD recognizes weak interspersed DNA homology as pairable

We engineered a series of synthetic Test DNA sequences, each related to Rsp1500 by a particular pattern of interspersed homology (Fig. 2E). These patterns were designed with an ‘**X**H-**Y**N’ format, in which homology tracts of **X** bp are separated by non-homology tracts of **Y** bp (Fig. 2E). The following patterns were analyzed: 7H-4N, 6H-5N, 5H-6N, 4H-7N, 3H-8N and 2H-1N (Fig. 2E). In addition, three instances of random homology were created using sequences obtained from *E. coli* and lambda phage. These sequences are unrelated to the ECS and LPS segments described above.

The presence of sensitizing heterology did not compromise the expected pairing of the identical Rsp1500 segments (compare Fig. 2D, ‘Identical alleles’ and Fig. 2F, ‘Perfect’). At the same time, all three instances of random homology produced 74-77% silenced ascospores (Fig. 2F, ‘Lambda’, ‘Ecoli1’, ‘Ecoli2’), substantially less than a deletion of the same length (Fig. 2D). This result suggests that the Rsp1500 segment on the first homologous chromosome becomes partially protected from MSUD by the presence of heterologous DNA at the allelic position on the second homologous chromosome. If the ‘Lambda’ DNA is extended by 3.5 kbp, the proportion of silenced ascospores increases to 87.7%, which corresponds to an intermediate level between random homology and the deletion (Fig. 2F, ‘Insertion’). This result suggests that the protection provided to Rsp1500 by heterologous allelic DNA is maximal when their lengths are matching.

Strikingly, interspersed homologies 7H-4N, 6H-5N and 5H-6N failed to induce MSUD, while 4H-7N yielded only a small (8.1%) fraction of silenced ascospores (Fig. 2F). Much stronger MSUD was triggered by 3H-8N (Fig. 2F). Finally, 2H-1N induced silencing at the level indistinguishable from that of random homology (Fig. 2F). To confirm that these effects were due to MSUD, we tried to suppress it by providing an ectopic 1500-bp fragment of *sad-2+* as the Test DNA. While this heterologous fragment should have activated strong MSUD, it was also expected to downregulate MSUD due to the ‘silencing the silencer’ effect. Indeed, only 13.4% of ascospores possessed the silenced phenotype in this case (while ∼75% were expected), suggesting that the observed effects were indeed caused by MSUD (Fig. 2F).

Taken together, our results show that weak interspersed homologies (characterized by an overall sequence identity below 50%, *e.g.*, patterns 4H-7N and 5H-6N) can be recognized by MSUD as pairable. These results also suggest that heterologous allelic sequences can escape MSUD, albeit less efficiently than homologous ones, with the maximal effect probably observed for matching sequence lengths.

### Recombination-independent recognition of DNA homology directs the expression of small RNAs

Previous studies identified a population of small RNAs (masiRNAs) produced specifically from the unpaired regions during MSUD (30). We investigated if the expression of these small RNAs was also regulated by the recognition of interspersed homologies. It was also important to ensure that the expression of masiRNAs in general is controlled by the process of DNA homology recognition rather than any unforeseen secondary effects, such as disruption of local chromatin structure by transformed DNA.

We started by asking if our Rsp1500 reporter system is suitable for pursuing this question. We analyzed two conditions expected to yield robust small RNA expression profiles. One condition featured the deletion, and the other one featured ‘Lambda’ DNA as an instance of random homology (Fig. 3A). Each of the unpaired segments produced small RNAs that mapped to both strands, and no spurious peaks were detected within the vicinity of the assayed regions (Fig. 3A). In general, the combined data demonstrate a good agreement between the genetic (ascospore eccentricity) and the molecular (small RNA-seq) readouts of MSUD using the Rsp1500 reporter (Fig. S5A,C).

**Figure 3.**
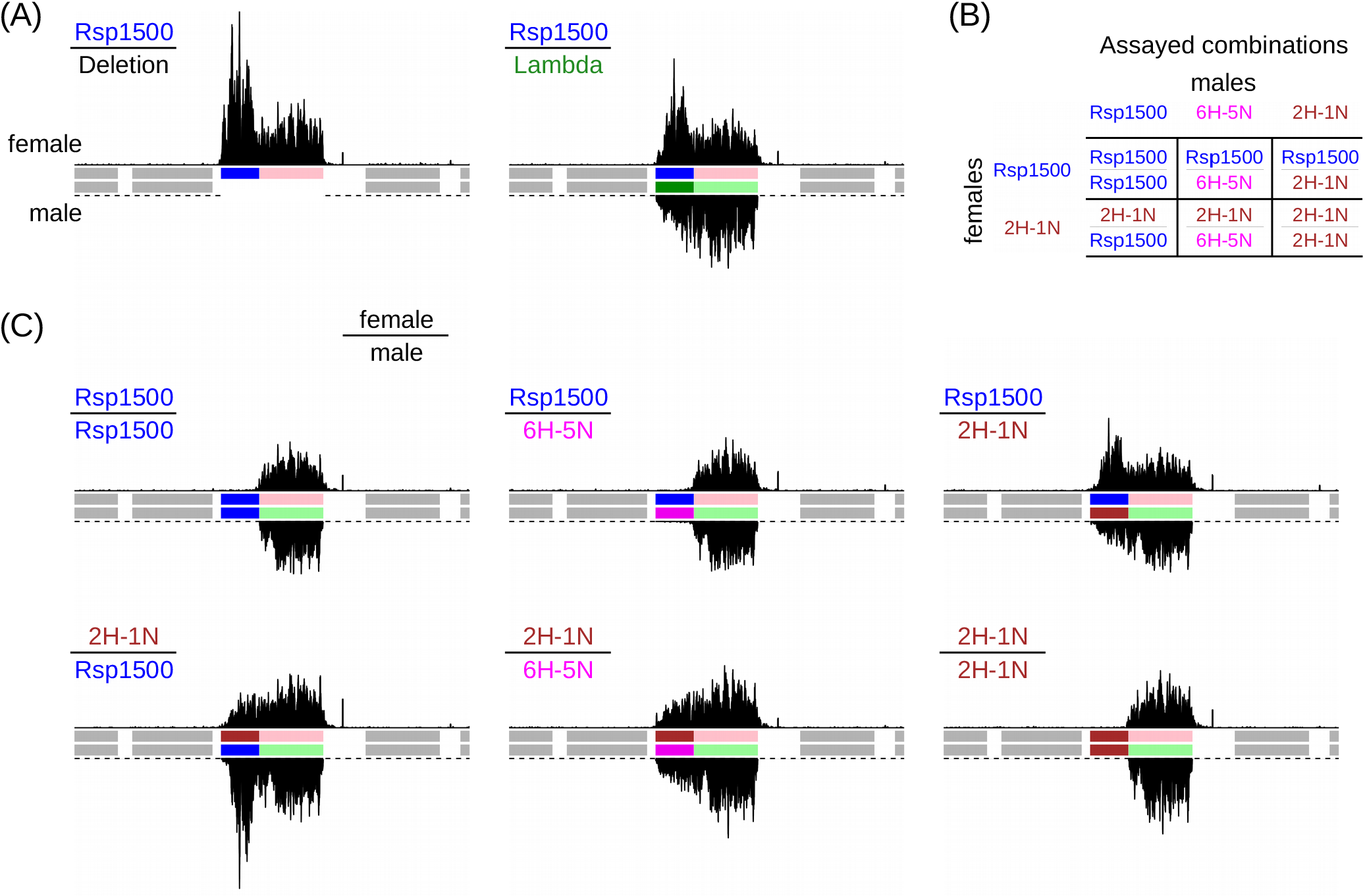
Recombination-independent recognition of DNA homology directs the expression of small RNAs. (A) Small RNA profiles. The following crosses are analyzed: X12 (’Deletion’) and X20 (’Lambda’) (SI Table 3). The identifiable species of small RNAs are limited to those carrying 5’ phosphate and 3’ hydroxyl (Methods). Neighboring genes are shown as gray rectangles. (B) The six combinations of the allelic 1500-bp segments. Female strains: RSP_Reporter1 (’Rsp1500’) and RSP_Reporter2 (’2H-1N’). Male strains: RSP_Test3 (’Rsp1500’), RSP_Test10 (’6H-5N’), and RSP_Test25 (’2H-1N’) (SI Table 2) (C) Small RNA profiles. The following crosses are analyzed: X13, X15, X19 (top row, left to right); X25, X26, and X27 (bottom row, left to right) (SI Table 3).

To perform a stringent test, an additional reporter strain (RSP_Reporter2) was created. Instead of Rsp1500 adjacent to ECS, this strain carried 2H-1N, also adjacent to ECS. RSP_Reporter1 and RSP_Reporter2 were separately crossed to three test strains carrying either Rsp1500, 6H-5N or 2H-1N. Therefore, these crosses featured six pairs of the 1500-bp segments, including two instances of perfect homology (Rsp1500/Rsp1500 and 2H-1N/2H-1N), three instances of ‘no homology’ (Rsp1500/2H-1N, 2H-1N/Rsp1500 and 2H-1N/6H-5N) and one instance of interspersed homology (Rsp1500/6H-5N) (Fig. 3B). The term ‘no homology’ refers to the fact that ‘Rsp1500/2H-1N’ induces *Rsp* silencing at the level of random homology (Fig. 2F).

Abundant small RNAs were expressed from all segments comprising the instances of ‘no homology’ (Fig. 3C). However, the same DNA constructs failed to express small RNAs above background in the context of perfect or interspersed homologies (Fig. 3C). These results demonstrate that the capacity to produce small RNAs is associated with the combined characteristics of two sequences rather than any individual sequence alone. Taken together, these results strongly suggest that small RNA expression during MSUD is mediated entirely by DNA homology recognition.

### Recognition of homology between Rsp1500 and Test DNA occurs within a larger genomic context

As reported previously, two identical DNA segments could be protected from MSUD even if they occupied somewhat different (non-co-linear) positions on a pair of homologous chromosomes (31). It was important to understand if this concept also applies when interspersed homologies are involved. To this end, we tested if partially homologous DNA segments remain protected from MSUD when they are inverted with respect to each other. Pattern 6H-5N was chosen for this analysis, because it is perceived as completely pairable with Rsp1500 in the direct (co-linear) orientation (Fig. 2F). As a control, we also examined the effect of inverting one of two Rsp1500 segments comprising perfect homology (Fig. S6A).

Strains carrying the inverted 6H-5N or Rsp1500 segments as Test DNAs were crossed to RSP_Reporter1. The non-co-linear Rsp1500 segments are still recognized as fully pairable (Fig. S6B), in agreement with the previous study (31). Surprisingly, inverting 6H-5N yields a large fraction of silenced ascospores (Fig. S6C). This effect is supported by small RNA-seq data (Fig. S6D). Nevertheless, the level of silencing induced by inverted 6H-5N is much lower than that induced by random homologies (53.6% versus ∼75%, respectively), suggesting that the detection of this pattern as pairable is strongly impeded but not completely abolished in the non-co-linear orientation. Taken together, these results show that recognition of homology between the allelic 1500-bp segments can be affected by their orientation with respect to each other and the surrounding genomic regions.

### Interspersed homologies with periods of 21 or 22 base-pairs are recognized as pairable

Thus far, our results show that interspersed homologies with the 11-bp periodicity are sensed by MSUD as pairable (Fig. 2F). To test if longer periodicities can also promote homology recognition, we engineered an additional series of 15 Test DNA sequences. Each Test DNA in this series contains 6-bp units of homology arrayed with a fixed periodicity ranging between 17 and 31 bp (Fig. 4A).

**Figure 4.**
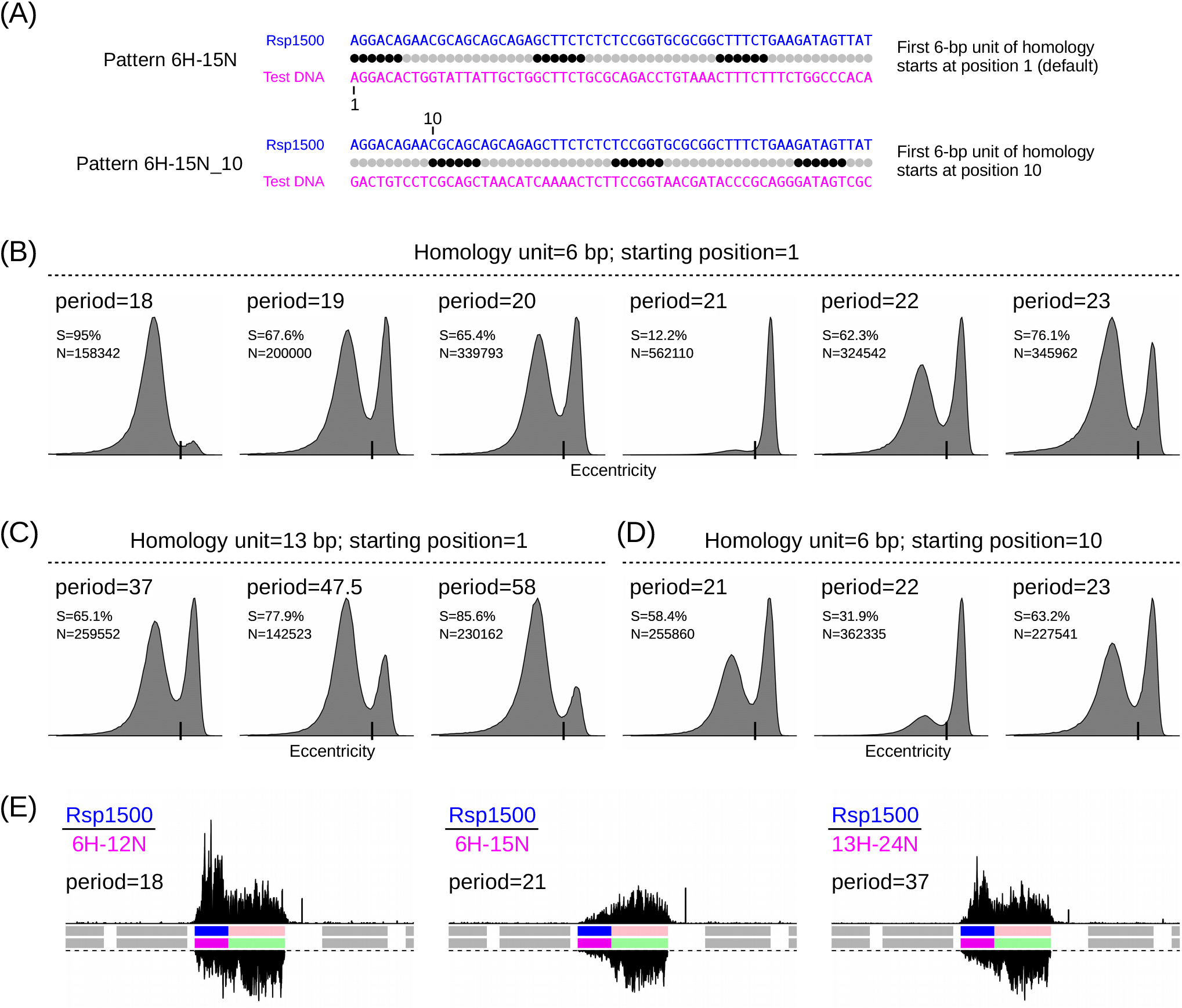
Interspersed homologies with periods of 21 or 22 base-pairs are recognized as pairable. (A) Two patterns 6N-15N that differ with respect to the starting positions of their homology frames. (B) Eccentricity distributions (displayed as in Fig. 2D) for the following crosses: X29, X30, X31, X32, X33, and X34 (left to right) (SI Table 3). (C) Eccentricity distributions (displayed as in Fig. 2D) for the following crosses: X43, X44, and X45 (left to right) (SI Table 3). (D) Eccentricity distributions (displayed as in Fig. 2D) for the following crosses: X46, X47, and X48 (left to right) (SI Table 3). (E) Small RNA profiles (displayed as in Fig. 3A) for the following crosses: X29, X32, and X43 (SI Table 3).

We found that pattern 6H-15N (period=21) yields a relatively small (12.2%) number of silenced ascospores, implying that it was often detected as paired. Patterns 6H-13N (period=19), 6H-14N (period=20) and 6H-16N (period=22) induced much higher levels of silencing (62─68%), and are therefore only marginally better than random homology at evading MSUD. Patterns 6H-11N (period=17) and 6H-12N (period=18), produced ∼95% silenced ascospores (Fig. 4B), a surprising outcome given that such strong MSUD response is typical of the deletion rather than any of our previous homology tests (Fig. 2). These results are fully consistent with the corresponding small RNA profiles (Fig. 4E). Lastly, the patterns with period lengths greater than 23 induced silencing at a level similar to random homology (Fig. S5A).

To probe the ability of 6H-15N to escape MSUD, we engineered additional variants of 6H-15N, 6H-16N and 6H-17N by changing the starting position of the 6-bp homology frames from 1 (default) to 10 (Fig. 4A). These new patterns were named 6H-15N_**10**, 6H-16N_**10**, and 6H-17N_**10**. Here we found that pattern 6H-15N_10 is considered less pairable than 6H-15N, while pattern 6H-16N_10 is considered more pairable than 6H-16N (Fig. 4B,D). At the same time pattern 6H-17N_10 is perceived as largely unpaired, which is similar to 6H-17N (Fig. 4B,D).

We also explored if some additional periods, expected to promote sequence-specific plectonemic pairing of relaxed B-DNAs (16), could be effective for the MSUD pairing process. These patterns contained 13-bp units of homology arrayed with the periodicity of 37, 47.5 (alternating 47-bp and 48-bp heterology tracts) or 58 bp (Fig. 4C). At first approximation, these patterns are recognized as unpaired (Fig. 4C). Taken together, these results reveal a distinct ability of the 21─22-bp periodicities to escape MSUD. However, this ability appears to be sequence-dependent, potentially reflecting the dynamic nature of the underlying homology detection process.

### An all-atom model of the direct homologous dsDNA-dsDNA pairing for MSUD

Our results reveal that MSUD is based on an efficient recombination-independent mechanism. Because this mechanism can detect weak interspersed homology (with an overall sequence identity level below 40%), it is unlikely to rely on single-stranded DNA or RNA intermediates. This mechanism is also precise, thus arguing against roles of the indirect sequence readouts. These and other (above) lines of evidence suggest that MSUD likely involves direct pairing of double-stranded DNA molecules.

An all-atom model of direct homologous dsDNA-dsDNA pairing has already been proposed (16). According to this model, one interaction unit contains 3-4 stacked quartets in the form of a short quadruplex. Each quartet is formed by two identical self-complementary base-pairs. Because of helical rotation constraints and concomitant structural deformations, two consecutive quadruplexes must be separated by more than one helical turn. For paranemic pairing, the spacing must be an integral number of helical turns, and it cannot be less than five (16). Yet for the right-handed plectonemic pairing, the minimal spacing distance (i) becomes shorter, (ii) depends upon the writhe of the plectoneme, and (iii) does not need to be an integer or half-integer in terms of free B-DNA helical turns (16). Thus, we investigated if the observed optimal 21─22-bp periodicity is compatible with such plectonemic pairing. To do so, we took a molecular dynamics (MD) approach, as explained below.

In principle, plectonemic structures were created as illustrated in Fig. S7A. Specifically, two parallel B-DNA double helices were attached to a virtual frame that consisted of a scaffolding pole and several crossbeams. This frame had only a few degrees of freedom corresponding to rotation of the crossbeams around the pole. It was built from phantom atoms not involved in any interactions except for harmonic links to some backbone groups. The frame was used in MD simulations for applying appropriate twisting torques that coiled the two straight helices into a plectoneme (Fig. S7A). The system was placed in a small water shell and neutralized by potassium ions (in vacuum). The MD simulations were carried out using established methods (32–34) that were previously applied to studying transitions between the A- and B-forms of DNA under low hydration (35–37).

The actual model was built in several steps (Fig. S7B). The initial conformation of the quadruplex structure was taken from one of the earlier MD trajectories (16). A complex with four stacked quartets was chosen and extended by canonical B-DNAs to the total length of 24 bp. Two such structures are shown in Fig. S7B (left panel) with orientations corresponding to the helical symmetry of a right-handed plectoneme. The lengths of the extended tails were 11 and 9 bp. In the final model, the shorter tails were ligated to obtain the spacing of 18 bp. This separation corresponds to interspersed homologies 3H-18N (period=21) or 4H-18N (period=22).

Next, the upper part was rotated and the red helices were ligated to create a structure shown in the second left panel (Fig. S7B). This structure was then attached to a virtual frame and covered by a hydration shell as explained above (Fig. S7A). Starting from this state, MD simulations were used to return the upper part back to its initial position. During this run, the red helix gradually assumed the shape corresponding to that in a right-handed plectoneme (second right panel in Fig. S7B). Once this step was complete, the free 8-bp ends of the blue helix were replaced by a continuous 16-bp fragment copied from the red helix. The accompanying structural perturbations and the rearranged hydration shell were re-equilibrated during several nanoseconds of MD simulations to produce the final structure (Fig. S7B, right panel).

Starting from the equilibrated structure (Fig. S7B, right panel), MD simulations were continued for several nanoseconds without any restraints, except having the DNA ends attached to the virtual frame to simulate the condition of long DNAs. The coordinates obtained during the last nanosecond were used for averaging (Fig. 5A). During this control run, DNA conformations fluctuated around the original state without significant local perturbations or general structural tendencies, suggesting that the paired structure is stable (Fig. 5B). The movie corresponding to the complete control run is provided as SI Data File 9.

**Figure 5.**
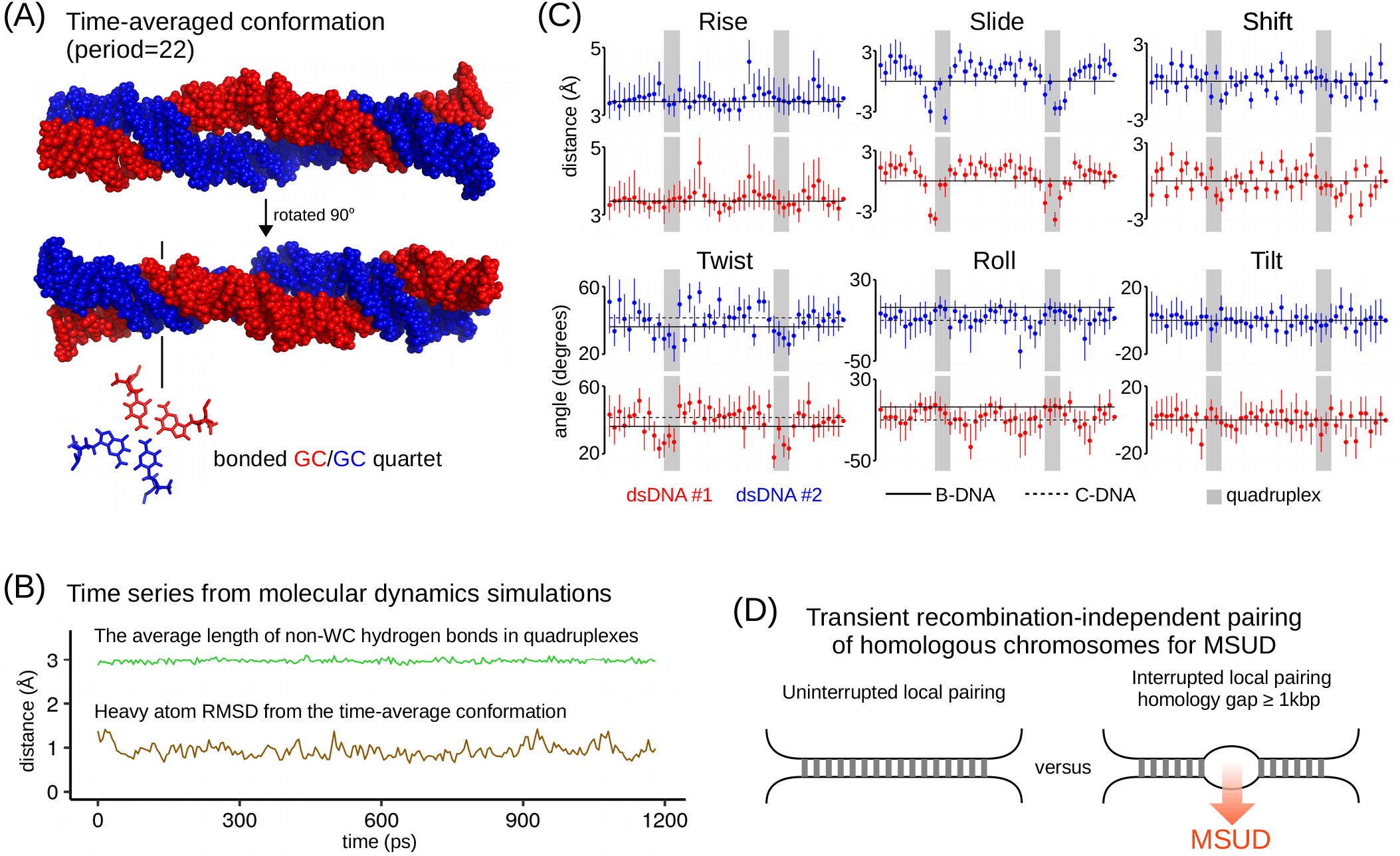
Direct dsDNA-dsDNA pairing as the basis for DNA homology recognition during MSUD. (A) The predicted structure of a paired complex with two DNA double helices of 48 bp. Two orthogonal views are shown of an average conformation sampled during one nanosecond of an all-atom molecular dynamics (MD) run with a shell of water and potassium ions (not shown). The spacing between the 4-bp quadruplexes corresponds to a period of 22 bp. The bonding of base-pairs inside the quadruplexes is illustrated for one quartet (not in scale). (B) Time traces of two geometric parameters that characterize the overall stability of the complex during the verification one-nanosecond MD run. (C) The principal helical parameters of the two dsDNA in the paired complex. MD averages are plotted, with error bars showing time fluctuations. The available experimental data (based on X-ray fiber diffraction) are shown by horizontal lines. The regions of quadruplex pairing are marked by gray shading. (D) The mechanism of triggering the production of small RNA in MSUD. Unpaired DNA can be recognized as being surrounded by the transiently paired regions.

Importantly, some helical parameters in the paired complex differed from the canonical B-DNA values (Fig. 5C). The most critical is the structural shift of unpaired segments towards the highly twisted C-DNA, which is typical of crystal fibers under low hydration conditions (38, 39). The uncovered B- to C-form transition is likely caused by the plectonemic coiling in combination with low hydration and high ion concentrations. Overall, the properties of the all-atom model (Fig. 5) indicate that the quadruplex-based pairing of dsDNA is, in principle, compatible with the optimal periodicities of 21─22 bp.

## DISCUSSION

Standard mechanisms of DNA homology recognition, including recombination (40), CRISPR/Cas9 (41) and DNA interference (42), use single-stranded DNA or RNA intermediates to probe dsDNA targets based on the Watson-Crick bonding principle. Consequently, if homology recognition needs to occur between two dsDNA molecules, one of them must be broken to create such an intermediate (43). Yet, paradoxically, homologous chromosomes can still pair in the absence of DNA breakage in many organisms, including *D. melanogaster* (46, 47), *C. elegans* (48), mice (44, 45) and fungi (49). In all of these situations, the specificity basis remains largely unknown.

The strongest evidence for a mechanism that matches intact dsDNAs is provided by Neurospora RIP, which can be induced by partial homologies with an overall sequence similarity of only 25─27%, provided that they contain short units of sequence identity interspersed with a periodicity of 11 or 12 bp (15). RIP occurs in the absence of MEI-3, the only RecA-like recombinase in *N. crassa* (15), thus setting itself further apart from the canonical recombinational processes.

In this study, we investigated the molecular requirements of meiotic silencing by unpaired DNA (MSUD) in *N. crassa*. During MSUD, any gene-sized chromosomal segment lacking a homologous allelic partner induces transient RNA interference in early meiosis (10, 11). We began by assaying the dependence of MSUD on the canonical mechanism of meiotic recombination, which requires SPO11 to cut dsDNAs and MEI-3 to catalyze the homology search (17). We found that both factors are completely dispensable for MSUD. We also found that SPO11-independent recombination (20) is not a relevant factor in our experimental system. This result was unexpected, because in *N. crassa* SPO11 is needed for robust chromosome synapsis (20). On the other hand, our findings hint that even when homolog synapsis in late prophase I requires SPO11 (*e.g.*, in budding yeast or mice), chromosomes are able to interact transiently by a recombination-independent mechanism prior to that. It is not impossible that such transient interactions are important for establishing subsequent synapsis.

Although MSUD does not need recombination for homology detection, it still faces a challenge of relying on interactions between homologous chromosomes rather than sister chromatids. Our results show that REC8, a universally conserved meiotic kleisin that promotes inter-chromosomal recombination, enhances MSUD. REC8 is normally loaded during premeiotic DNA replication (23), which takes place before karyogamy in *N. crassa* and related fungi (50). We found that MSUD is somewhat retarded in the absence of REC8, but it still occurs in most meiotic nuclei. The recombination-independent role for REC8 in MSUD may be related to its function in favoring general inter-chromosomal interactions during meiosis.

A possibility of MSUD being independent from meiotic recombination was discussed earlier (29), yet its basis remained mysterious, in part because the properties of MSUD appeared compatible with a strand annealing mechanism (29). Specifically, it was found that a pair of allelic sequences with only 6% divergence (produced by RIP) could trigger MSUD (29). Given the homology requirements of RIP (15), we investigated if MSUD could also recognize interspersed homologies with sequence divergence greater than 50% as pairable. We created a sensitive genetic assay to accurately measure the response of MSUD to interactions between two 1500-bp allelic segments. The sequence of one (reporter) segment remained always the same, whereas the other (test) segment was varied as desired. A similar approach was used earlier in our studies of RIP (15).

By systematically testing different patterns of homology, we found that MSUD shares its homology-sensing principles with RIP. Specifically, MSUD treats short tracts of sequence identity interspersed with the 11-bp periodicity as pairable. Like in RIP, the capacity of MSUD to recognize such interspersed homologies can be affected by the adjacent regions of perfect homology. Some pairable homologies in our assays corresponded to an overall sequence identity of only 36.4%, thus arguing against the role of strand annealing at the basis of homology recognition during MSUD. On the other hand, the assayed interspersed homologies did not fully mimic perfect homology (for example, by the co-linearity test, Fig. S6), pointing to the existence of additional sequence requirements that were not accommodated by the designed patterns. For example, the homology search may proceed in several steps (*e.g.*, coarse pairing followed by fine pairing) with distinct requirements that can be fulfilled by perfect homology, but not the interspersed homologies examined in our assays.

By systematically exploring interspersed homologies with periods ranging between 17 and 31, we found that periods of 21─22 bp could support pairing. In our system, the exact value of the optimal periodicity depends on which frame of homologous units is chosen. Thus, pairing is influenced by position and sequence effects. We used the 22-bp periodicity as a constraint in all-atom modeling of plectonemic DNA structures linked by interspersed quadruplexes (16). Our effort yielded a paired dsDNA-dsDNA complex that is consistent with standard chemical geometry and that does not interfere with intrahelical WC bonding (according to empirical potentials currently used in molecular dynamics simulations of DNA). The complex contains 4-bp quadruplex nodes connected by plectonemically coiled dsDNA segments. The conformation of these ‘free’ segments is similar to that of C-DNA. The strongly twisted double helix of C-DNA derives from B-DNA by increasing the fraction of phosphate groups switched from the canonical BI to an alternative BII backbone conformation (51). The minor groove of C-DNA is deep and narrow, and the major groove of C-DNA is wide and relatively flat. Thus, the quadruplex-based recognition and pairing of homologous C-DNAs can be very fast, satisfying the speed requirements of RIP and MSUD. The B-to-C DNA transition likely depends on ion concentrations and molecular crowding in the immediate vicinity of the participating DNAs.

Our results suggest that homologous chromosomes engage in transient recombination-independent pairing in early prophase I. Such pairing may feature sequence-specific quadruplex-based contacts along with non-specific interactions. The ability of the heterologous allelic segments to escape MSUD at an appreciable rate suggests that non-specific interactions may play an important role. Overall, if the homologous chromosomal regions do not contain a large (< 1 kbp) section of heterology, they will associate more or less continuously; however, if such heterology is present, it will remain largely unpaired and will be recognized as such in the context of the paired flanking regions (Fig. 5D).

The proposed pairing mechanism likely relies on many additional factors, notably those involved in chromatin remodeling. Indeed, one nucleosome remodeler, SAD-6 (a SWI/SNF helicase), has already been implicated in MSUD (31). In yeast, a considerable fraction of histones is removed by a proteasome-mediated process in response to DNA damage to facilitate recombinational DNA repair (52); and proteasome-mediated protein turnover has been implicated in homologous pairing and recombination during meiosis (53, 54). It is possible that these pathways are also relevant for recombination-independent homologous dsDNA-dsDNA pairing.

## METHODS

### DNA constructs

All constructs were created as plasmids using standard molecular cloning techniques. Synthetic DNAs were designed as previously described (15) and ordered from Integrated DNA Technologies as ‘gBlocks’. All 1500-bp DNA fragments assayed in this study are provided in FASTA format in SI Data File 1. Natural sequences were amplified from commercially available sources using Phusion polymerase (NEB, cat. no. M0530S). The *hH1-gfp* gene was obtained from pMF280 (19). All plasmids used in this study are listed in SI Table 1. Maps of all these plasmids are provided in EMBL format in SI Data File 2.

### Manipulation of Neurospora strains

All Neurospora strains used in this study are listed in SI Table 2. Standard media were used for growth and crosses. Vogel’s Medium N with 1.5% sucrose was used for unrestricted vegetative growth. Colonial growth was induced by plating macroconidia or ascospores on sorbose agar (3% agar, 2% sorbose, 0.1% dextrose, 1x Vogel’s Medium N salts). For transformation, ∼1-2 μg of linearized plasmid DNA was electroporated into macroconidia in the total volume of 25 μl using the following parameters: 1500 V, 600 Ω, 25 μF, 2 mm gap. Homokaryotic strains were purified from primary heterokaryotic transformants by macroconidiation. Inserts were validated by sequencing. Crosses were setup in 90-mm Petri dishes (Gosselin, cat. no SB90-101) on 1x Synthetic Crossing medium with 2% sucrose. Nylon filtration fabric with pore size 250 μm was used for support. Crosses were started by placing ∼10^3^ macroconidia (in 1-2 μl of water) at the center of each plate. On the fifth day after inoculation, mycelial lawns were fertilized using suspensions of macroconidia of the opposite mating type. Plates were then kept in the dark at 24°C in a Heratherm incubator (model IMP180). Ejected ascospores started to appear on the lids 8-9 days after fertilization. Original lids were replaced on day-10 after fertilization. The production of ascospore continued for 1-2 weeks. Ascospores were collected from the lids in drops of water on day-24 after fertilization. All crosses analyzed in this study are listed in SI Table 3.

### Assaying hH1-GFP expression

Perithecia were fixed in a solution of 100 mM PIPES (pH 6.9), 10 mM EGTA, 5 mM MgSO_4_ and 4% PFA for 20 minutes at room temperature as previously described (55). Perithecia were washed and stored in sodium phosphate buffer (80 mM Na_2_HPO_4_, 20 mM NaH_2_PO_4_). Asci were dissected in 25% glycerol on glass support and transferred into a drop of mounting medium (25% glycerol, 10 mg/mL DABCO, 5 µg/mL Hoechst 33342, and 100 mM potassium phosphate buffer at pH 8.7). Cover slips were placed over samples and sealed with a thin layer of nail polish. Slides were stored at −20°C, if needed. Images were acquired using a 20× objective on a Zeiss Axio Imager M2 equipped with a PIXIS 1024 CCD camera (Princeton Instruments). Contrast and color were adjusted in Fiji with ‘Image>Adjust>Brightness/Contrast’ and ‘Image>Lookup Tables’, respectively (56). Separate images are also provided in SI Data File 3.

### Assaying recombination

Ascospores were collected from 24-day crosses, heat-shocked in water (60°C for 30 min) and plated directly on sorbose agar containing cyclosporin A (5 μg/ml; Sigma, cat. no. 30024). Genomic DNA was extracted as previously described (15). A combination of four primers was used to type the *mat* locus by PCR. One pair (5’-AAGTATCGCCAAAGCTGGTTC-3’ and 5’-TCATGGCAAAGTCCAACTTCC-3’) amplified *mat A*, while the other pair (5’-TCCCGGACTTCACAATAACGA-3’ and 5’-GCGCGAAGTTTTCTAGATCCT-3’) amplified *mat a*. One negative control (’no DNA’) and two positive controls (both parental strains) were included in each set of 30 reactions prepared from the same master mix. PCR products were resolved on 1× TAE 1% agarose gels and imaged using Gel Doc EZ System (Bio-Rad). Raw gel scans are provided in SI Data File 4. The scans can be viewed with Fiji (56).

### Assaying *Roundspore* phenotype: collecting images

Freshly collected suspensions of ascospores were concentrated to approximately 10^5^ per ml. Glass-bottom 35-mm dishes (Ibidi, cat. no. 81156) were used for imaging. Dishes were pre-loaded with 0.4 ml of water and mounted on a motorized stage of a Zeiss Axio Observer Z1. Suspensions of ascospores (0.1 ml) were added and mixed thoroughly by pipetting. Once ascospores ceased to sediment, a 16×16 array of non-overlapping image tiles was acquired using Hamamatsu ORCA-Flash4.0 V3 camera (C13440). The following parameters were used: PlnN 10×/0.3 DICI objective, 1.6× Optovar and DIC illumination. Several 16×16 arrays could be acquired per cross. JPEG images (2048×2048 pixels) were exported using ZEN software (Zeiss). Tiles overlapping the edges of the glass window were removed manually.

### Assaying *Roundspore* phenotype: segmenting and classifying ascospores

Ascospores were detected by a two-step image-processing algorithm, in which the rough object localization was followed by the local segmentation. The rough localization relied on the detection of ascospore edges. First, noise was removed by applying a median filter. To gain computational speed, the image was first down-scaled by a factor of two. A median filter with a radius of five pixels was used, after which the image was up-scaled back to its original size. A gradient filter with a fixed radius of two pixels was then applied to highlight the ascospore edges, and a binary mask was generated by thresholding that filtered image. Rough contours were thinned to obtain a line contour, which was then expanded by one pixel. Roughly segmented objects corresponded to the pixels inside those contours. The objects were filtered based on their area values, and the retained regions were subjected to local segmentation. Small subimages centered on the detected regions (bounding box ± 10 pixels) were extracted and automatically thresholded. The objects were filtered to retain those above the convexity threshold of 0.97. Their roundness was approximated by the eccentricity parameter, which could vary between 0 (circle) and 1 (linear segment). Data from a complete set of images (obtained from three replica crosses) were combined to calculate the total fraction of ‘silenced’ ascospores. The algorithm is written in Python and available at github.com/guiwitz/sporeseg. Version v0.0.1 was used. The parameters of all analyzed ascospores are provided as plain-text tables in CSV format in SI Data File 5.

### Preparing and sequencing small RNA libraries

Total RNA was extracted from 6-day-old perithecia with TRI Reagent (Sigma-Aldrich, cat. no. T9424) and re-suspended in 5M urea. RNA with a RINe value above 8 (as determined using Agilent TapeStation 2200) was fractionated on a 8% TBE-Urea gel to selected RNA molecules in the 17-26 nt range. Excised gel slices were crushed and incubated overnight in 0.3 M NaCl at 25°C while shaking at 850 rpm. Gel debris was separated from RNA-containing solution using 0.22-µm spin columns (Sigma-Aldrich, cat. no. CLS8161). Small RNAs were precipitated with four volumes of 100% ethanol and centrifugation at 20000 ×g (for 40 minutes at 4°C). RNA pellets were re-suspended in 4 µl of RNA-free water (per tube, per two gel lanes, corresponding to 40-80 µg of total RNA). Purified RNAs were used to construct libraries as previously described (57), except that all cleaning steps used Agencourt RNAClean XP Beads (Beckman, cat. no. A63987). Libraries were purified on 6% non-denaturing polyacrylamide gels, quantified on TapeStation 2200 and sequenced on NextSeq 500 (Illumina).

### Small RNA sequence analysis

Reads were demultiplexed with bcl2fastq v2.20 (Illumina), trimmed with cutadapt v1.15 (58), and aligned to custom meta-assemblies with bowtie2 v2.3 (59). Each custom meta-assembly corresponded to a unique pair of parental strains. Meta-assemblies were based on the reference NC12 (GCF_000182925.2) and included the following additional contigs: transformed DNAs, mitochondrial DNA (NC_026614.1) and the *mat a* locus (M54787.1). The 1500-bp segment of the endogenous *Rsp+* gene corresponding to Rsp1500 was masked with N’s. If transformed DNA was shared by the parents, it was masked with N’s in the ‘male’ contig. A meta-assembly corresponding to a cross between RSP_Reporter1 and the test strain carrying 6H-5N is provided as an example (in EMBL format, with all the contigs merged into one sequence) in SI Data File 6.

SAM and BAM files were manipulated with samtools v1.7 (60). All reads that mapped to the ribosomal locus were removed. To depict small RNA profiles, a normalization procedure was implemented. Specifically, 148 regions with strong and consistent expression of small RNAs were annotated manually based on the entire RNA-seq dataset. These regions ranged from 1000 to 28900 bp in length (816.6 kbp in total) and accounted for 60-70% of all nonribosomal small RNAs mapped to the nuclear genome. The sequences of these regions are provided in FASTA format in SI Data File 7. The position of each region in the provided meta-assembly is indicated in the header. The total number of reads and the number of reads mapped to the standard regions are listed in SI Table 4. Corresponding BAM files are provided in SI Data File 8. Profiles were visualized with ggplot2 (61).

### Model building and molecular simulations

All molecular simulations involved in model building were performed with internal coordinate method (ICMD) (62, 63) using the standard fixed geometry of chemical groups and the internal mobility of DNA limited to essential degrees of freedom. In contrast, equilibration and stability verification of the final state were carried out with all degrees of freedom of DNA using conventional MD in Cartesian coordinates. The DNA duplexes were modeled using a recent version of the AMBER force field (64, 65) in a small shell of SPC/E water (66) neutralized by potassium ions (67) under vacuum boundary conditions. The electrostatic interactions were treated by the custom version of the smooth particle mesh Ewald (SPME) method (68) originally developed for MD simulations of water drops (32). Additional details and references concerning the simulation protocols can be found elsewhere (69). The movie corresponding to the last nanosecond is provided in MP4 format in SI Data File 9.

## SUPPLEMENTARY TABLES

**SI Table 1.** Plasmids used in this study.

| Plasmid ID | Assayed DNA ID | Assayed DNA source | ECS/LPS | Reference |
| --- | --- | --- | --- | --- |
| pEAG54 | Deletion | N/A | N/A | this study |
| pEAG258D | hH1-GFP | pMF280 (ref. 19) | N/A | this study |
| pEAG267A | hH1-GFP | pMF280 (ref. 19) | N/A | this study |
| pTSN3A | Insertion | NC_001416.1: 10130-16322 | N/A | this study |
| pTSN4B | Identical | NC_026501.1: 9283898-9282399 | ECS | this study |
| pTSN4D | Lambda | NC_001416.1: 12636-14135 | LPS | this study |
| pTSN4E | Perfect | NC_026501.1: 9283898-9282399 | LPS | this study |
| pTSN4F | Perfect(inverted) | NC_026501.1: 9283898-9282399 | LPS | this study |
| pTSN4H | 13H-24N | Synthetic | LPS | this study |
| pTSN4I | 13H-34.5N | Synthetic | LPS | this study |
| pTSN4J | 13H-45N | Synthetic | LPS | this study |
| pTSN4K | 6H-5N | Synthetic | LPS | this study |
| pTSN4L | 6H-5N(inverted) | Synthetic | LPS | this study |
| pTSN4M | 6H-15N | Synthetic | LPS | this study |
| pTSN4N | 6H-16N | Synthetic | LPS | this study |
| pTSN4O | 6H-17N | Synthetic | LPS | this study |
| pTSN4P | 6H-18N | Synthetic | LPS | this study |
| pTSN4Q | 6H-19N | Synthetic | LPS | this study |
| pTSN4R | 6H-11N | Synthetic | LPS | this study |
| pTSN4S | 6H-12N | Synthetic | LPS | this study |
| pTSN4T | 6H-13N | Synthetic | LPS | this study |
| pTSN4U | 6H-14N | Synthetic | LPS | this study |
| pTSN4V | 7H-4N | Synthetic | LPS | this study |
| pTSN4W | 5H-6N | Synthetic | LPS | this study |
| pTSN4X | 4H-7N | Synthetic | LPS | this study |
| pTSN4Y | 3H-8N | Synthetic | LPS | this study |
| pTSN9B | 2H-1N | Synthetic | LPS | this study |
| pTSN9C | 6H-15N_10 | Synthetic | LPS | this study |
| pTSN9F | 6H-16N_10 | Synthetic | LPS | this study |
| pTSN9G | 6H-17N_10 | Synthetic | LPS | this study |
| pTSN9H | 6H-23N | Synthetic | LPS | this study |
| pTSN9I | 6H-24N | Synthetic | LPS | this study |
| pTSN9J | Ecoli2 | NC_010473.1: 1725001-1726500 | LPS | this study |
| pTSN9K | Ecoli1 | NC_010473.1: 3744001-3745500 | LPS | this study |
| pTSN9N | 6H-20N | Synthetic | LPS | this study |
| pTSN9P | Sad-2 | NC_026505.1: 5531277-5532776 | LPS | this study |
| pTSN9R | 2H-1N | Synthetic | ECS | this study |
| pTSN9S | 6H-21N | Synthetic | LPS | this study |
| pTSN9T | 6H-22N | Synthetic | LPS | this study |
| pTSN9U | 6H-25N | Synthetic | LPS | this study |

ECS: NC\_010473.1: 1636254-1638753
LPS: NC\_001416.1: 10130-12629

**SI Table 2.** Neurospora strains used in this study. To avoid RIP, all strains listed below had both *rid+* and *dim-2Δ+* deleted.

| Strain ID | Other ID | Mat | Genotype | Construction strategy | Reference |
| --- | --- | --- | --- | --- | --- |
| GFP_Report1 | C194.3 | a | *hH1-gfp::mei-3Δ, spo11Δ, csr-1Δ* | crossing, *hH1-gfp* from pEAG258D | this study |
| REC_Test1 | C50.3 | A | *his-3* | published previously | ref. 22 |
| REC_Test2 | C245.5 | A | *sad-2Δ* | crossing | this study |
| REC_Test3 | RNR136.9 | A | *mei-3Δ, spo11Δ* | crossing | this study |
| REC_Test4 | C289.4 | A | *mei-3Δ, spo11Δ, sad-2Δ* | crossing | this study |
| REC_Test5 | C196.4 | A | *hH1-gfp::mei-3Δ, spo11Δ* | crossing, *hH1-gfp* from pEAG258D | this study |
| REC_Test6 | C192.10 | A | *spo11Δ* | crossing | this study |
| GFP_Report2 | T568.1 | A | *rec8Δ, spo11Δ, hH1-gfp::csr-1Δ* | crossing, *hH1-gfp* from pEAG267A | this study |
| REC8_Test1 | C237.1 | a | *rec8Δ, spo11Δ,* | crossing | this study |
| REC8_Test2 | C244.18 | a | - | crossing | this study |
| GFP_Report3 | T583.1 | A | *dim-5Δ, set-7Δ, hH1-gfp::csr-1Δ* | crossing, *hH1-gfp* from pEAG267A | this study |
| HET_Test1 | C252.9 | a | *dim-5Δ, set-7Δ* | crossing | this study |
| HET_Test2 | C252.4 | a | *dim-5Δ, set-7Δ, sad-2Δ* | crossing | this study |
| C58.7 | C58.7 | a | *his-3* | published previously | ref. 22 |
| RSP_Report1 | C254.36 | A | *Rsp1500::ECS* | cross progeny of T571 and C50.3 | this study |
| RSP_Report2 | C284.32 | A | *2H-1N::ECS* | cross progeny of T642 and C50.3 | this study |
| RSP_Test1 | T571 | a | *Rsp1500::ECS* | C58.7 transformed with pTSN4B | this study |
| RSP_Test2 | T575 | a | *Lambda::LPS* | C58.7 transformed with pTSN4D | this study |
| RSP_Test3 | T576 | a | *Rsp1500::LPS* | C58.7 transformed with pTSN4E | this study |
| RSP_Test4 | T577 | a | *Rsp1500(inverted)::LPS* | C58.7 transformed with pTSN4F | this study |
| RSP_Test5 | T579 | a | *13H-24N::LPS* | C58.7 transformed with pTSN4H | this study |
| RSP_Test6 | T580 | a | *13H-34.5N::LPS* | C58.7 transformed with pTSN4I | this study |
| RSP_Test7 | T581 | a | *13H-45N::LPS* | C58.7 transformed with pTSN4J | this study |
| RSP_Test8 | T582 | a | *Insertion* | C58.7 transformed with pTSN3A | this study |
| RSP_Test9 | T585 | a | *Deletion* | C58.7 transformed with pEAG54 | this study |
| RSP_Test10 | T586 | a | *6H-5N::LPS* | C58.7 transformed with pTSN4K | this study |
| RSP_Test11 | T587 | a | *6H-5N(inverted)::LPS* | C58.7 transformed with pTSN4L | this study |
| RSP_Test12 | T588 | a | *6H-15N::LPS* | C58.7 transformed with pTSN4M | this study |
| RSP_Test13 | T589 | a | *6H-16N::LPS* | C58.7 transformed with pTSN4N | this study |
| RSP_Test14 | T590 | a | *6H-17N::LPS* | C58.7 transformed with pTSN4O | this study |
| RSP_Test15 | T591 | a | *6H-18N::LPS* | C58.7 transformed with pTSN4P | this study |
| RSP_Test16 | T592 | a | *6H-19N::LPS* | C58.7 transformed with pTSN4Q | this study |
| RSP_Test17 | T602 | a | *6H-11N::LPS* | C58.7 transformed with pTSN4R | this study |
| RSP_Test18 | T603 | a | *6H-12N::LPS* | C58.7 transformed with pTSN4S | this study |
| RSP_Test19 | T604 | a | *6H-13N::LPS* | C58.7 transformed with pTSN4T | this study |
| RSP_Test20 | T605 | a | *6H-14N::LPS* | C58.7 transformed with pTSN4U | this study |
| RSP_Test21 | T606 | a | *7H-4N::LPS* | C58.7 transformed with pTSN4V | this study |
| RSP_Test22 | T607 | a | *5H-6N::LPS* | C58.7 transformed with pTSN4W | this study |
| RSP_Test23 | T608 | a | *4H-7N::LPS* | C58.7 transformed with pTSN4X | this study |
| RSP_Test24 | T609 | a | *3H-8N::LPS* | C58.7 transformed with pTSN4Y | this study |
| RSP_Test25 | T612 | a | *2H-1N::LPS* | C58.7 transformed with pTSN9B | this study |
| RSP_Test26 | T613 | a | *6H-15N_10::LPS* | C58.7 transformed with pTSN9C | this study |
| RSP_Test27 | T625 | a | *6H-16N_10::LPS* | C58.7 transformed with pTSN9F | this study |
| RSP_Test28 | T626 | a | *6H-17N_10::LPS* | C58.7 transformed with pTSN9G | this study |
| RSP_Test29 | T627 | a | *6H-23N::LPS* | C58.7 transformed with pTSN9H | this study |
| RSP_Test30 | T628 | a | *6H-24N::LPS* | C58.7 transformed with pTSN9I | this study |
| RSP_Test31 | T629 | a | *Ecoli2::LPS* | C58.7 transformed with pTSN9J | this study |
| RSP_Test32 | T630 | a | *Ecoli1::LPS* | C58.7 transformed with pTSN9K | this study |
| RSP_Test33 | T640 | a | *Sad-2::LPS* | C58.7 transformed with pTSN9P | this study |
| RSP_Test34 | T642 | a | *2H-1N::ECS* | C58.7 transformed with pTSN9R | this study |
| RSP_Test35 | T652 | a | *6H-20N::LPS* | C58.7 transformed with pTSN9N | this study |
| RSP_Test36 | T653 | a | *6H-21N::LPS* | C58.7 transformed with pTSN9S | this study |
| RSP_Test37 | T654 | a | *6H-22N::LPS* | C58.7 transformed with pTSN9T | this study |
| RSP_Test38 | T655 | a | *6H-25N::LPS* | C58.7 transformed with pTSN9U | this study |

**SI Table 3.** Crosses analyzed in this study.

| Cross ID | Female ID | Female other ID | Male ID | Male other ID |
| --- | --- | --- | --- | --- |
| X1 | REC_Test1 | C50.3 | GFP_Reporter1 | C194.3 |
| X2 | REC_Test2 | C245.5 | GFP_Reporter1 | C194.3 |
| X3 | REC_Test3 | RNR136.9 | GFP_Reporter1 | C194.3 |
| X4 | REC_Test4 | C289.4 | GFP_Reporter1 | C194.3 |
| X5 | REC_Test5 | C196.4 | GFP_Reporter1 | C194.3 |
| X6 | REC_Test6 | C192.10 | GFP_Reporter1 | C194.3 |
| X7 | REC8_Test1 | C237.1 | GFP_Reporter2 | T568.1 |
| X8 | REC8_Test2 | C244.18 | GFP_Reporter2 | T568.1 |
| X9 | HET_Test1 | C252.9 | GFP_Reporter3 | T583.1 |
| X10 | HET_Test2 | C252.4 | GFP_Reporter3 | T583.1 |
| X11 | RSP_Reporter1 | C254.36 | RSP_Test1 | T571 |
| X12 | RSP_Reporter1 | C254.36 | RSP_Test9 | T585 |
| X13 | RSP_Reporter1 | C254.36 | RSP_Test3 | T576 |
| X14 | RSP_Reporter1 | C254.36 | RSP_Test21 | T606 |
| X15 | RSP_Reporter1 | C254.36 | RSP_Test10 | T586 |
| X16 | RSP_Reporter1 | C254.36 | RSP_Test22 | T607 |
| X17 | RSP_Reporter1 | C254.36 | RSP_Test23 | T608 |
| X18 | RSP_Reporter1 | C254.36 | RSP_Test24 | T609 |
| X19 | RSP_Reporter1 | C254.36 | RSP_Test25 | T612 |
| X20 | RSP_Reporter1 | C254.36 | RSP_Test2 | T575 |
| X21 | RSP_Reporter1 | C254.36 | RSP_Test32 | T630 |
| X22 | RSP_Reporter1 | C254.36 | RSP_Test31 | T629 |
| X23 | RSP_Reporter1 | C254.36 | RSP_Test33 | T640 |
| X24 | RSP_Reporter1 | C254.36 | RSP_Test8 | T582 |
| X25 | RSP_Reporter2 | C284.32 | RSP_Test3 | T576 |
| X26 | RSP_Reporter2 | C284.32 | RSP_Test10 | T586 |
| X27 | RSP_Reporter2 | C284.32 | RSP_Test25 | T612 |
| X28 | RSP_Reporter1 | C254.36 | RSP_Test17 | T602 |
| X29 | RSP_Reporter1 | C254.36 | RSP_Test18 | T603 |
| X30 | RSP_Reporter1 | C254.36 | RSP_Test19 | T604 |
| X31 | RSP_Reporter1 | C254.36 | RSP_Test20 | T605 |
| X32 | RSP_Reporter1 | C254.36 | RSP_Test12 | T588 |
| X33 | RSP_Reporter1 | C254.36 | RSP_Test13 | T589 |
| X34 | RSP_Reporter1 | C254.36 | RSP_Test14 | T590 |
| X35 | RSP_Reporter1 | C254.36 | RSP_Test15 | T591 |
| X36 | RSP_Reporter1 | C254.36 | RSP_Test16 | T592 |
| X37 | RSP_Reporter1 | C254.36 | RSP_Test35 | T652 |
| X38 | RSP_Reporter1 | C254.36 | RSP_Test36 | T653 |
| X39 | RSP_Reporter1 | C254.36 | RSP_Test37 | T654 |
| X40 | RSP_Reporter1 | C254.36 | RSP_Test29 | T627 |
| X41 | RSP_Reporter1 | C254.36 | RSP_Test30 | T628 |
| X42 | RSP_Reporter1 | C254.36 | RSP_Test38 | T655 |
| X43 | RSP_Reporter1 | C254.36 | RSP_Test5 | T579 |
| X44 | RSP_Reporter1 | C254.36 | RSP_Test6 | T580 |
| X45 | RSP_Reporter1 | C254.36 | RSP_Test7 | T581 |
| X46 | RSP_Reporter1 | C254.36 | RSP_Test26 | T613 |
| X47 | RSP_Reporter1 | C254.36 | RSP_Test27 | T625 |
| X48 | RSP_Reporter1 | C254.36 | RSP_Test28 | T626 |
| X49 | RSP_Reporter1 | C254.36 | RSP_Test4 | T577 |
| X50 | RSP_Reporter1 | C254.36 | RSP_Test11 | T587 |

**SI Table 4.** Small RNA libraries analyzed in this study.

| Female<br>other ID | Male<br>other ID | Small RNA<br>lib ID | Mapped<br>reads % | Ribosomal<br>reads | MitoDNA<br>reads | Nuclear<br>genome | Standard<br>regions | Standard<br>regions (%) |
| --- | --- | --- | --- | --- | --- | --- | --- | --- |
| C254.36 | T585 | RRS30_1 | 96.32 | 847346 | 47492 | 3279975 | 2216383 | 67.57 |
| C254.36 | T575 | RRS9_0 | 97.05 | 1275847 | 58428 | 5555623 | 3893338 | 70.08 |
| C254.36 | T576 | RRS10_2 | 96.46 | 923966 | 50617 | 3586703 | 2391358 | 66.67 |
| C254.36 | T586 | RRS14_1 | 97.12 | 762401 | 54357 | 2974659 | 2105525 | 70.78 |
| C254.36 | T612 | RRS13_0 | 97.05 | 1572347 | 136121 | 5714123 | 4053507 | 70.94 |
| C284.32 | T576 | RRS19_1 | 95.84 | 636785 | 44907 | 3623259 | 2362889 | 65.21 |
| C284.32 | T586 | RRS21_1 | 97.02 | 650308 | 36581 | 3921834 | 2708514 | 69.06 |
| C284.32 | T612 | RRS22_1 | 96.94 | 756155 | 44985 | 4170856 | 2752629 | 66.00 |
| C254.36 | T603 | RRS26_1 | 95.55 | 622211 | 68486 | 1998732 | 1298578 | 64.97 |
| C254.36 | T588 | RRS23_1 | 94.64 | 1350357 | 123450 | 3192414 | 1952108 | 61.15 |
| C254.36 | T579 | RRS31_0 | 97.35 | 1572510 | 108027 | 5402130 | 3719496 | 68.85 |
| C254.36 | T587 | RRS15_2 | 97.14 | 991440 | 78289 | 3538723 | 2514906 | 71.07 |

## SUPPLEMENTARY DATA FILES

All data files are available as TAR/GZIP archives at figshare.com/s/6669fb33f3b249a41b25

## ACKNOWLEDGMENTS

We thank Sarah Bellout, Amy Boyd, Tyler Malone, Pennapa Manitchotpisit, Turner Reid, Pegan Sauls, and Aykhan Yusifov for technical assistance during early stages of this project. The work was supported by the grants from Fondation pour la Recherche Médicale (AJE20180539525), Agence Nationale de la Recherche (10-LABX-0062, 11-LABX-0011, ANR-19-CE12-0002), the National Institutes of Health (1R15HD076309-01), Institut Pasteur, and CNRS.

## AUTHOR CONTRIBUTIONS

E.G. and T.H. proposed the study; N.R., T.N., A.K.M, and E.G performed experiments; G.C. advised on small RNA sequencing and analysis; G.W. created the image processing algorithm; E.G. wrote the manuscript, to which A.K.M. and T.H. provided critical inputs.

---

**Figure S1.**
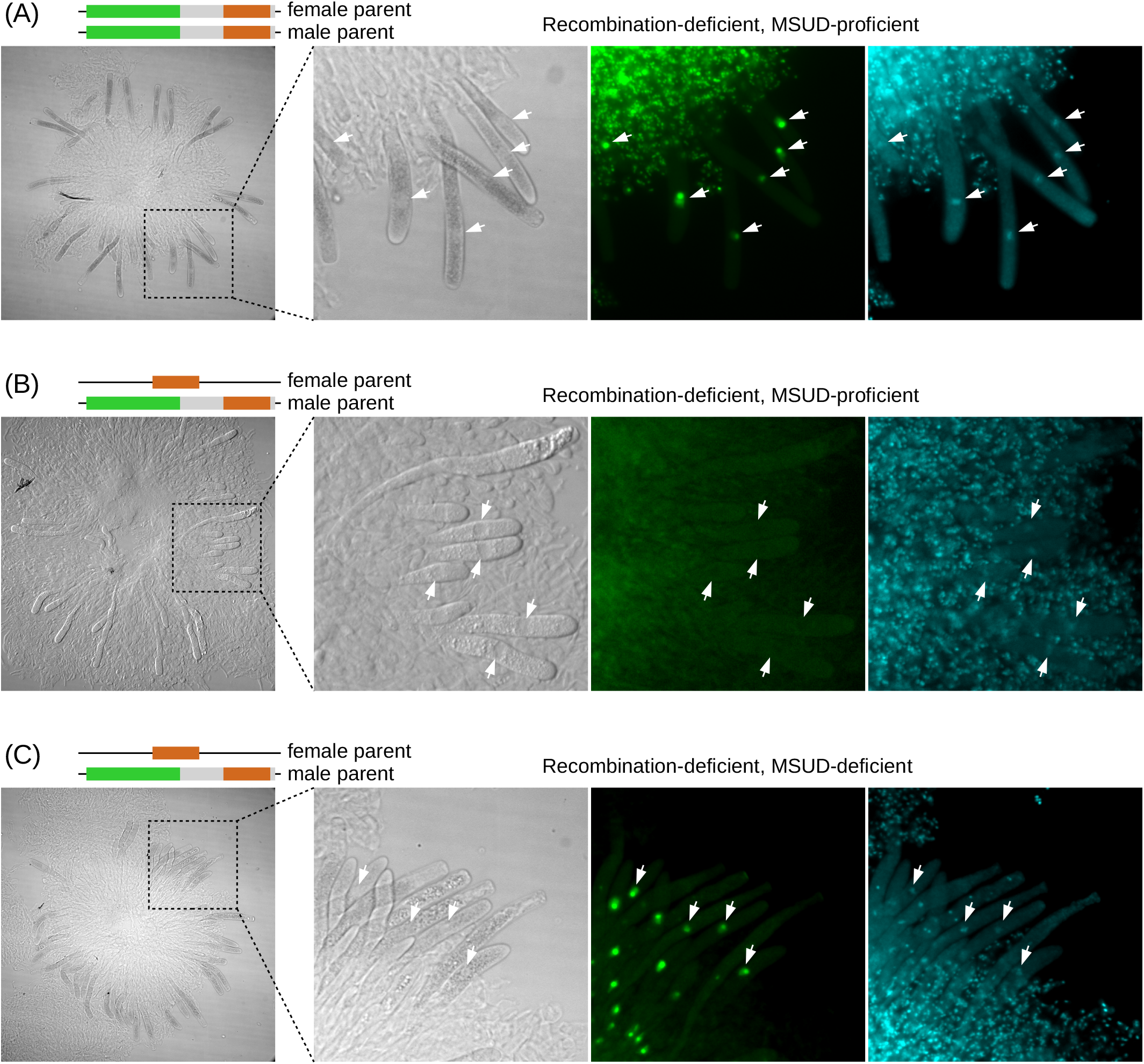
Pairable *hH1-gfp* alleles are expressed in the recombination-deficient background. (A) Cross X5 (SI Table 3). (B) Cross X3 (SI Table 3). (C) Cross X4 (SI Table 3). Representative meiotic nuclei are indicated with white arrows. Right panels show Hoechst staining.

**Figure S2.**
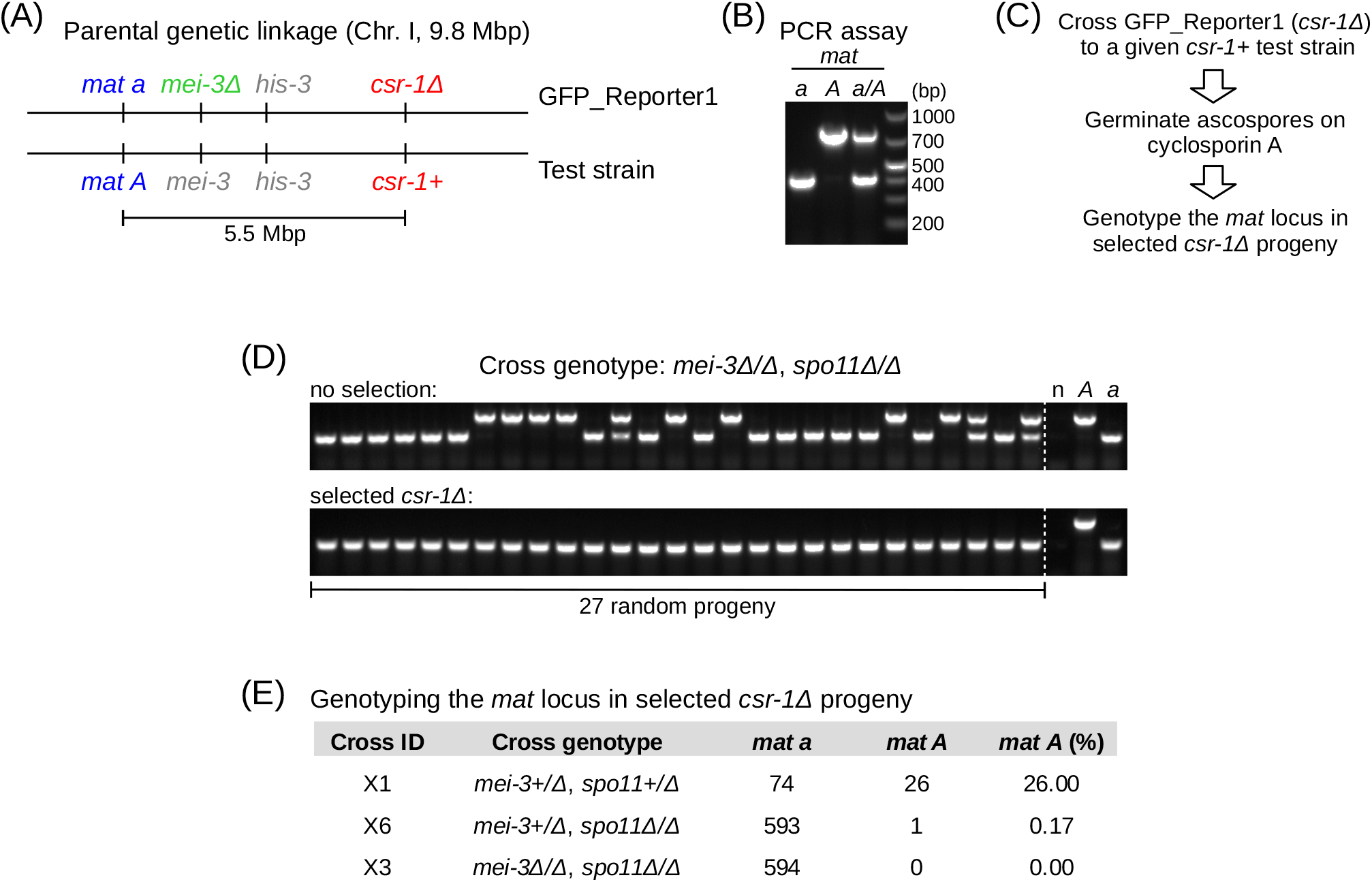
MSUD is not coupled to genetic recombination. (A) Genetic recombination is measured between the *mat* and *csr-1* loci, over more than half of chromosome I. This interval includes the *his-3* locus reported to undergo SPO11-independent recombination (20). (B) An approach to detect the concomitant presence of *mat a* and *mat A* by multiplex PCR. (C) GFP_Reporter1 (*mat a*, *csr-1Δ*) is crossed to test strains carrying *mat A* and *csr-1+*. (D) A sample of 27 random progeny from the recombination-deficient cross X3 (SI Table 3) were germinated in the absence (top) and presence (bottom) of cyclosporin A. (E) The number of *mat a* and *mat A* progeny obtained in three different genetic conditions, as indicated.

**Figure S3.**
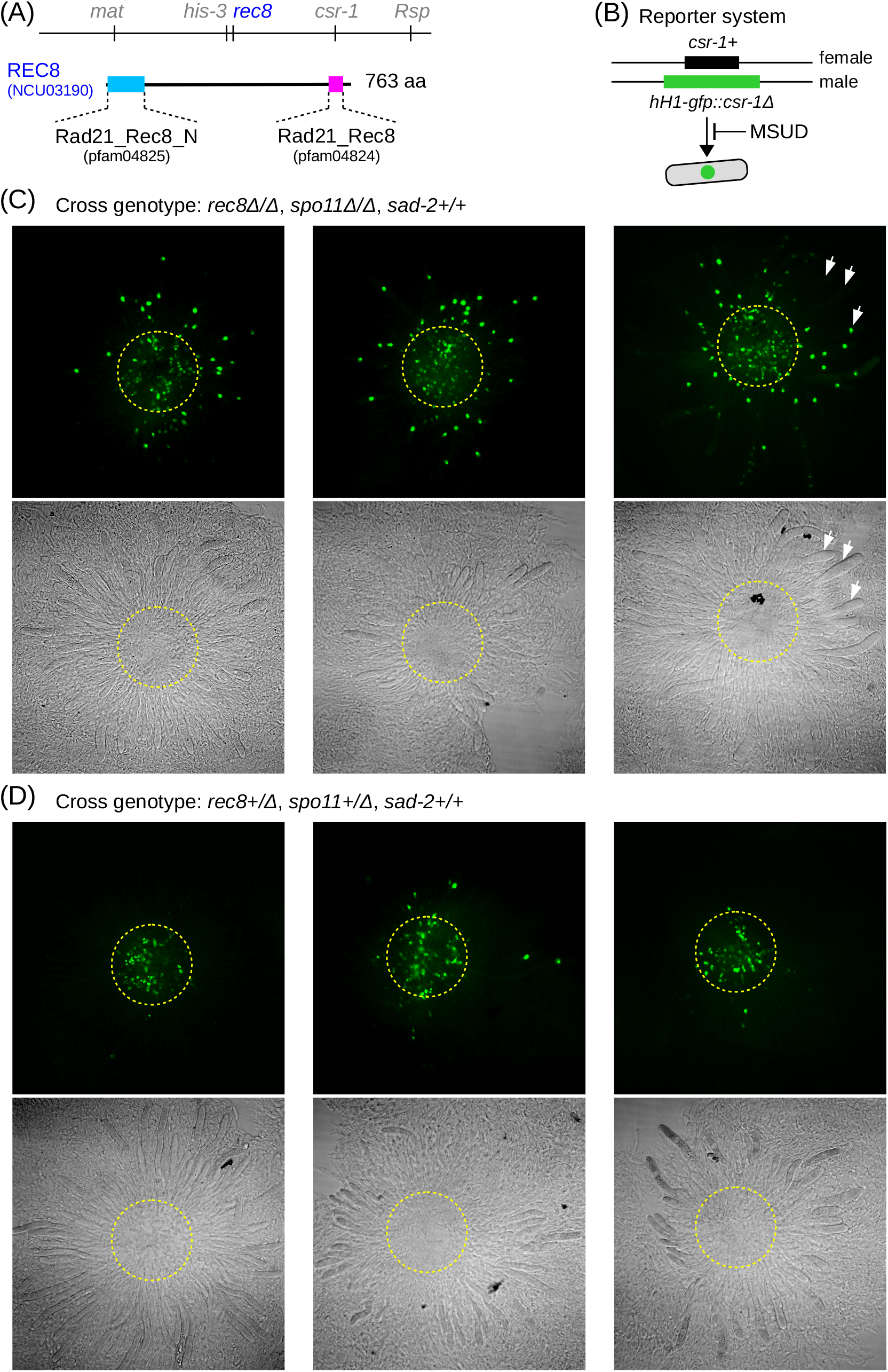
MSUD is promoted by REC8. (A) REC8 is encoded by NCU03190 and has a canonical structure comprising the conserved N-terminal and C-terminal domains. (B) The *hH1-gfp* construct is integrated as the replacement of the wildtype *csr-1+* allele. (C) Three representative rosettes from the REC8-deficient cross X7 (SI Table 3). (D) Three representative rosettes from the REC8-proficient cross X8 (SI Table 3). The approximate area containing premeiotic and very early meiotic nuclei is outlined (yellow dotted circles). Representative meiotic nuclei are indicated with white arrows.

**Figure S4.**
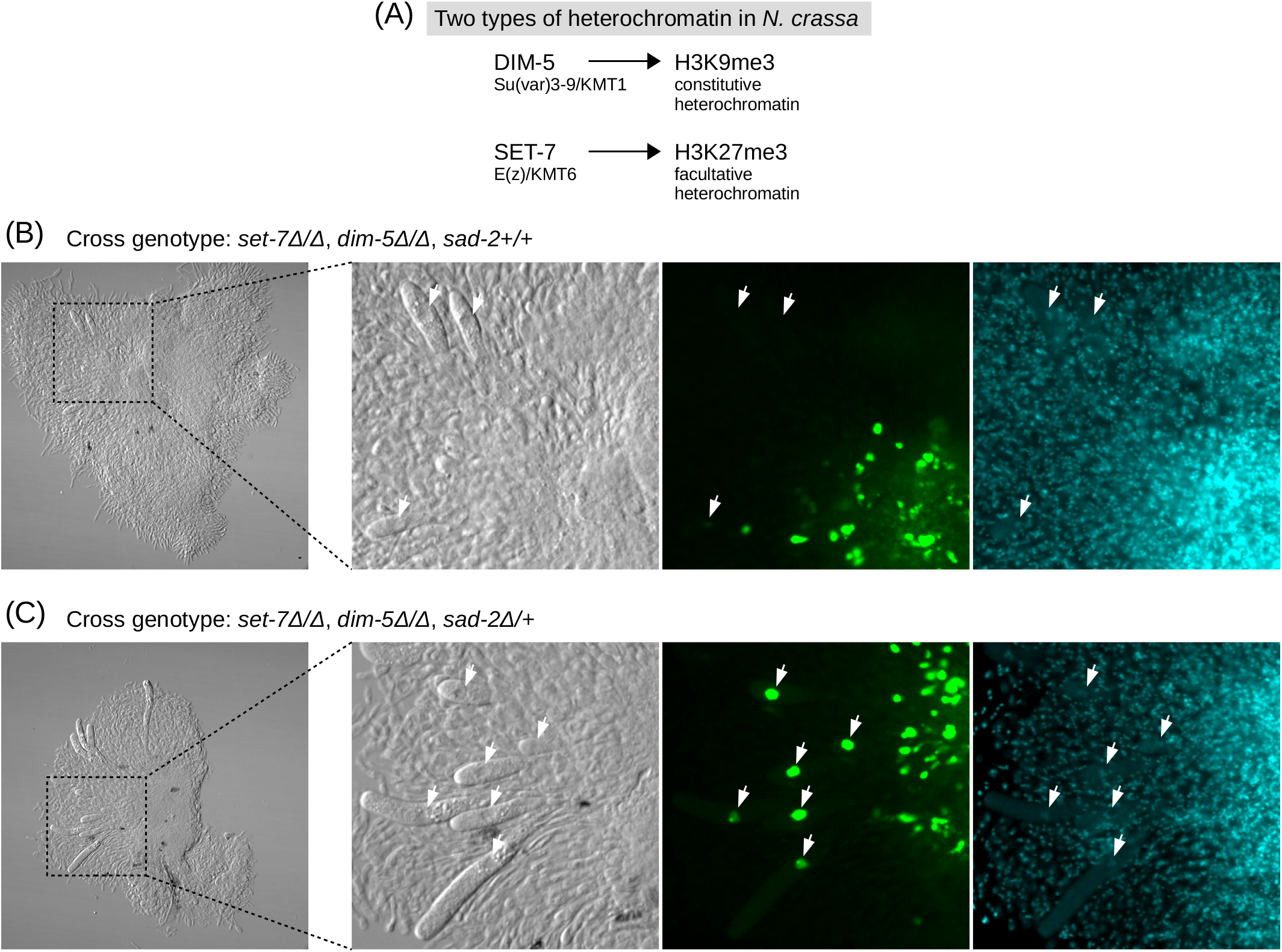
MSUD does not depend on the canonical mechanisms of heterochromatin formation. (A) *N. crassa* possesses two types of heterochromatin. Constitutive heterochromatin features H3K9me3 mediated by DIM-5, facultative heterochromatin features H3K27me3 mediated by SET-7. (B) Heterochromatin-deficient, MSUD-proficient cross X9 (SI Table 3). (C) Heterochromatin- and MSUD-deficient cross X10 (SI Table 3). Representative meiotic nuclei are indicated with white arrows. Right panels show Hoechst staining.

**Figure S5.**
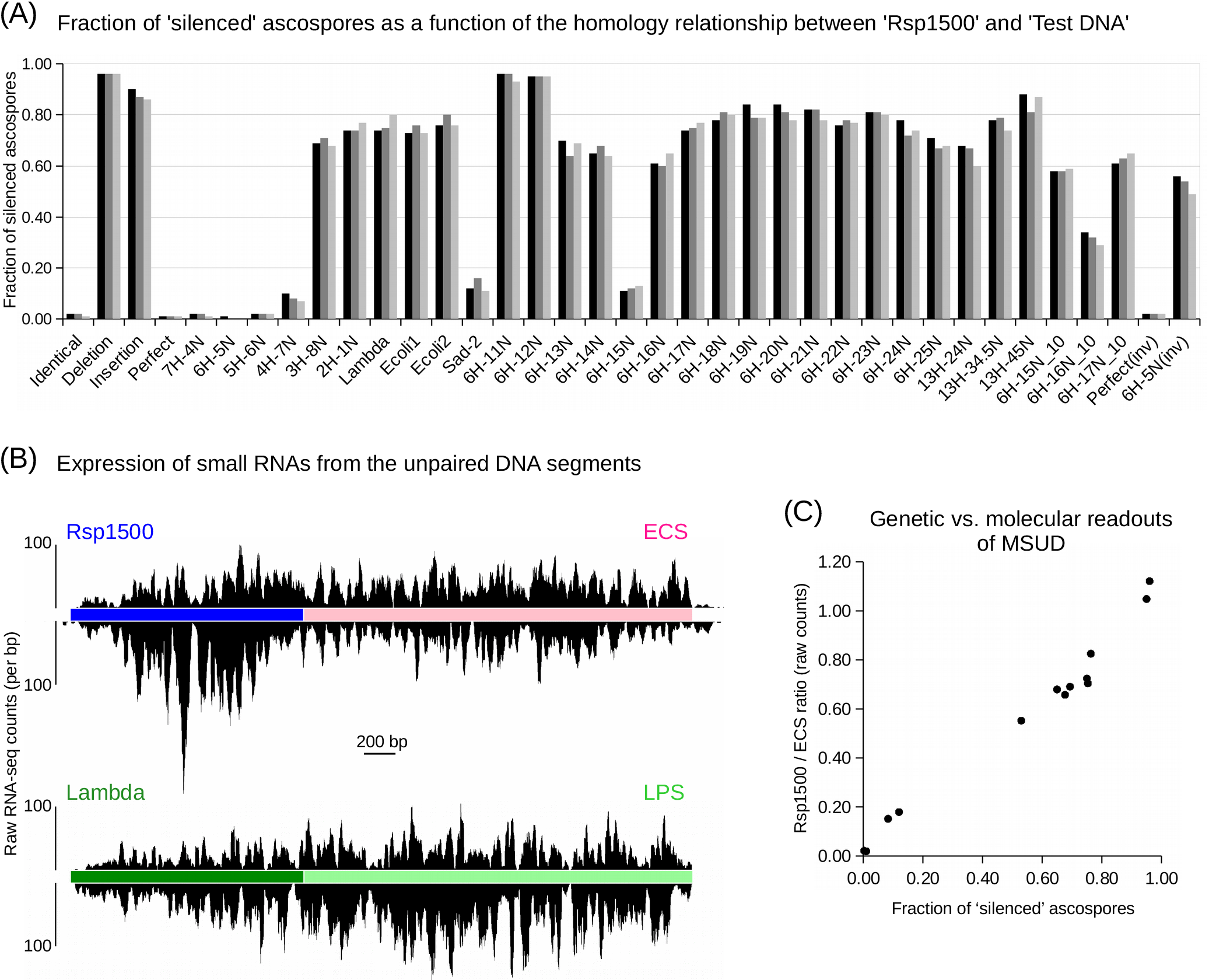
Properties of the Rsp1500 reporter system. (A) The proportion of silenced ascospores for individual replica crosses (in three different shades of gray). RSP_Reporter1 is used as a standard female parent. Except for Test DNAs, all male strains are isogenic. (B) Strand-specific expression of small RNAs from the unpaired DNA regions in cross X20 (SI Table 3). (C) The relationship between the fraction of silenced ascospores and the ratio of the total number of small RNA reads mapped to the ‘Rsp1500’ and ‘ECS’ segments of RSP_Reporter1 (SI Table 3).

**Figure S6.**
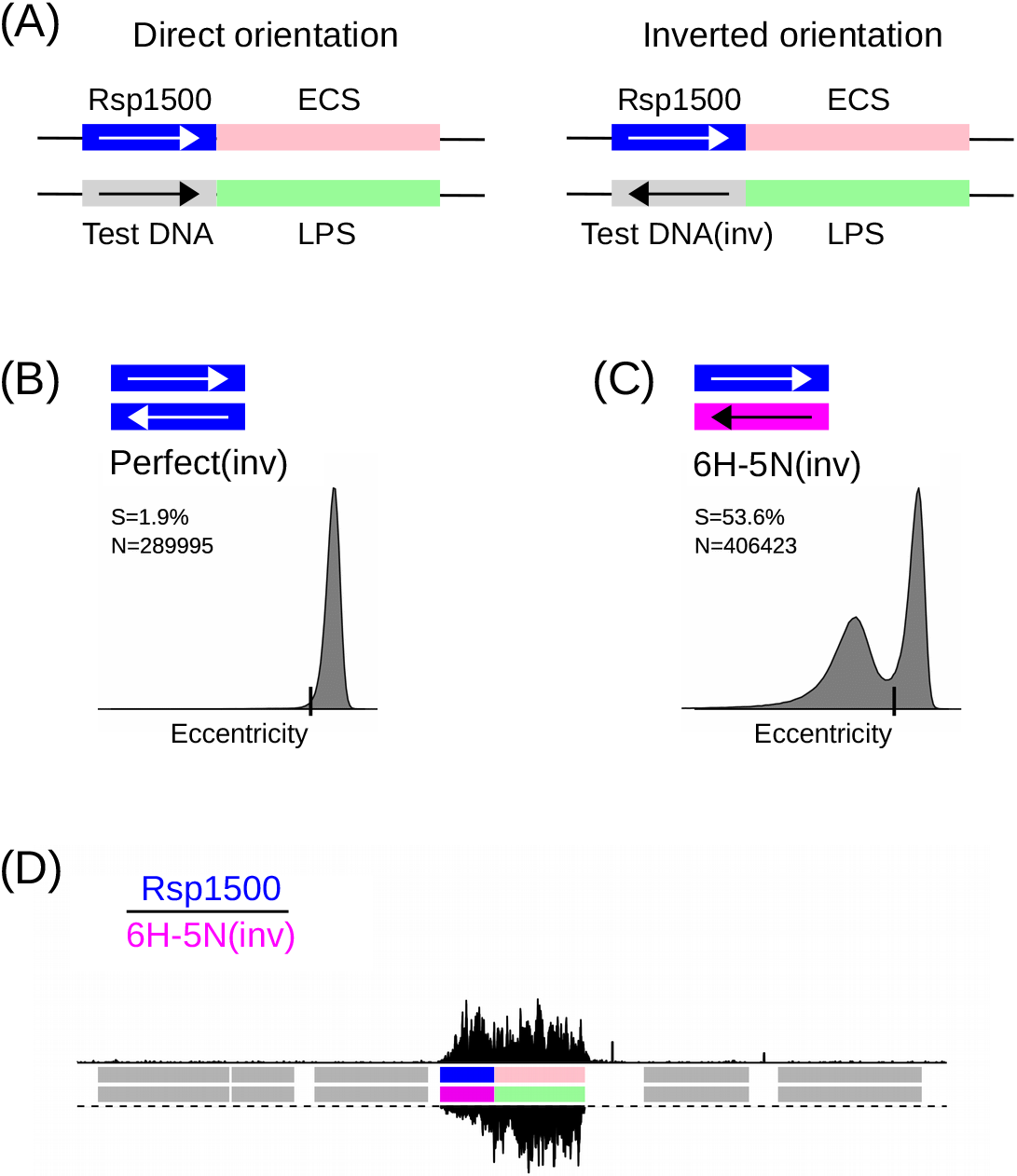
Recognition of homology between Rsp1500 and Test DNA occurs within a larger genomic context. (A) Two non-co-linear situations are examined involving either inverted perfect homology or inverted interspersed homology 6H-5N. (B) Eccentricity distribution (displayed as in Fig. 2D) corresponding to the inverted perfect homology. Cross X49 (SI Table 3). (C) Eccentricity distribution (displayed as in Fig. 2D) corresponding to inverted homology 6H-5N. Cross X50 (SI Table 3). (D) Small RNA profile (displayed as in Fig. 3A) corresponding to the eccentricity distribution in panel C.

**Figure S7.**
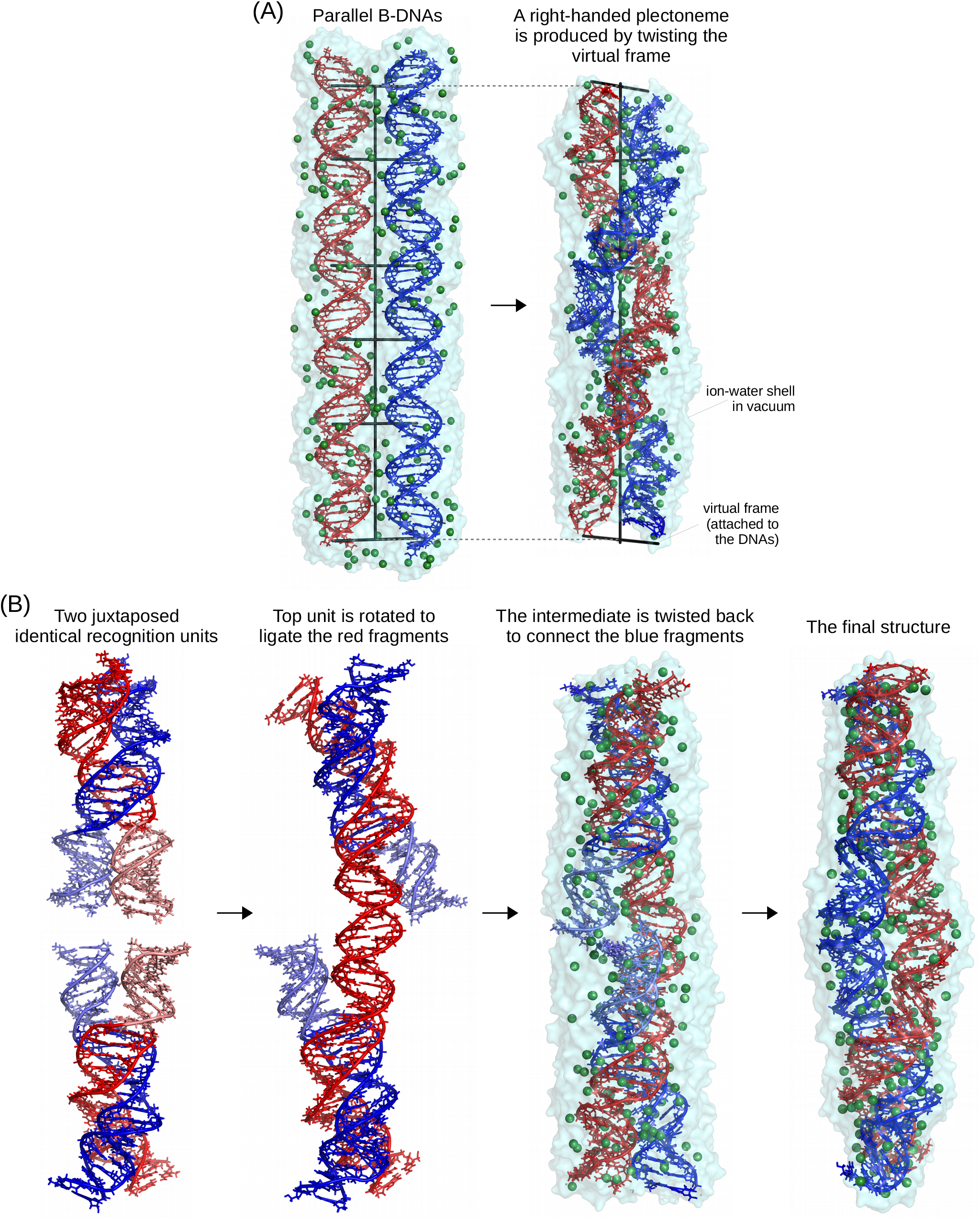
Constructing an all-atom model of homologous dsDNA-dsDNA pairing. (A) The method used to smoothly fold two straight double helices into a right-handed plectoneme. (B) The main steps of assembling a complex containing two paired 48-bp DNAs. The details are explained in the main text.

